# Fragmentation through cell-cell adhesion maintains senescent cell viability but promotes debris deposition

**DOI:** 10.1101/2023.01.11.523642

**Authors:** Matej Durik, Daniel Sampaio Gonçalves, Tania Knauer-Meyer, Coralie Spiegelhalter, Nadia Messaddeq, Marco Seehawer, Dmitry Bulavin, Lars Zender, William M. Keyes

**Affiliations:** Institut de Génétique et de Biologie Moléculaire et Cellulaire (IGBMC), Équipe Labellisée Ligue Contre le Cancer, Illkirch, France; UMR7104, Centre National de la Recherche Scientifique (CNRS), Illkirch, France; U1258, Institut National de la Santé et de la Recherche Médicale (INSERM), Illkirch, France; Université de Strasbourg, Illkirch, France; Department of Internal Medicine VIII, University Hospital Tübingen, 72076 Tübingen, Germany; Department of Physiology I, Institute of Physiology, Eberhard Karls University Tübingen, 72076 Tübingen, Germany; Translational Gastrointestinal Oncology Group, German Consortium for Translational Cancer Research (DKTK), German Cancer Research Center (DKFZ), Heidelberg 69120, Germany; Institute for Research on Cancer and Aging of Nice (IRCAN), Université Côte d’Azur, INSERM, CNRS, Nice, France

**Author notes:** Correspondence to ***Bill Keyes,*** IGBMC, 1 Rue Laurent Fries, BP 10142, 67404 Illkirch - CU Strasbourg, France, ***Matej Durik***, IGBMC, 1 Rue Laurent Fries, BP 10142, 67404 Illkirch - CU Strasbourg, France.

**Keywords:** senescence, mitochondria, cell-cell adhesion, debris, aging, cancer, DAMPs

## Abstract

Cellular senescence is a state of stable arrest and secretion linked to aging and disease. Here we identify that senescent cells dispose of large fragments of themselves through cell-to-cell adhesion, which we term senescent-cell adhesion fragments (SCAFs). Found across all senescent types examined, SCAFs lack nuclear material but contain organelles, including damaged mitochondria. Disrupting adherens junctions decreased SCAF formation but induced senescent-cell death, which was caused by an inability to shed damaged mitochondria. Dynamic analyses show that SCAFs ultimately rupture, releasing a complex proteome including damage-associated molecular patterns (DAMPs) and proteins linked to neurodegenerative disease. Functionally, SCAFs activate wound-healing and cancer-related programs, promoting migration and invasion. Altogether, these findings identify a new feature that facilitates senescent cell survival, but the consequence of which is external deposition of damaged intracellular contents, with implications for cancer and aging.

## INTRODUCTION

Cellular senescence is a highly dynamic and complex cellular response that can affect all stages of life.^1,2^ Senescence is triggered in response to replicative exhaustion, oncogene activation, chemotherapy and various cellular stresses, but it can also be induced in a controlled manner in the context of regeneration or embryonic development.^2–5^ Two main characteristics of senescent cells are an irreversible proliferative arrest and a highly active secretory profile called the senescence-associated secretory phenotype (SASP).^4,6^ The SASP consists of many signaling molecules such as chemokines, growth factors and extracellular matrix remodelers, and promotes the recruitment of immune cells to remove senescent cells.^7– 9^

The outcome of senescence induction can either be beneficial or detrimental, depending on timing and context. Senescent cells play physiological roles during embryonic development, wound healing and regeneration, where they appear transiently, and through the SASP, contribute to tissue patterning, regeneration, growth and repair.^2,10–12^ However, it is thought that this process becomes misregulated during aging, with the aberrant accumulation of senescent cells actively contributing to tissue aging.^13,14^ In addition, many age-related diseases have an increased senescent cell burden, ranging from idiopathic pulmonary fibrosis (IPF) and arthritis, to Alzheimer’s disease (AD), Amyotrophic Lateral Sclerosis (ALS) and Parkinson’s disease.^3,15–17^ Senescent cells also significantly impact cancer formation, promoting tumor cell proliferation, recurrence after treatment and metastasis.^5,18^ Despite frequently containing significant macromolecular damage, including damaged organelles, senescent cells remain alive, and are even resistant to cell death. During recent years, numerous studies have identified how the elimination or removal of senescent cells in aged or diseased states can be beneficial.^13–15,19,20^ Therefore, aberrant induction of senescence is detrimental, and interfering with it can have a beneficial impact on aging and age-related diseases.

However, exactly how cellular senescence contributes to aging or pathology remains unresolved. There is a critical need to better understand the cellular mechanisms of senescence, how these cells persist despite substantial damage, and how they contribute to aging and disease. Here, we unexpectedly find that senescent cells dispose of large fragments of themselves through cell–cell adhesion. We show that this fragmentation is required for senescent-cell survival, yet ultimately results in the rupture of these fragments and the release of diverse intracellular cargo onto neighboring cells and into the extracellular space. We propose that senescent fragments represent another critical step towards our understanding of the link with senescent cells and aging and disease processes.

## RESULTS

### Senescent cells deposit large fragments through cell-cell adhesion

Cellular senescence is a complex process, with typical markers, including senescence-associated beta-galactosidase (SA-β-gal) activity, and p21 and p16 expression appearing dynamically after induction (Extended data Fig. 1). Our interest was to study the dynamic process of senescence onset and maintenance. To achieve this, we established an *in vitro* model system that would enable us to image and track individual cells becoming senescent over time (Fig. 1A). This involved infecting human IMR-90 fibroblasts with a constitutive GFP-expressing retroviral vector (MSCV-PIG), so that after drug selection, these cells would stably express GFP. Using these GFP-positive fibroblasts, we then either induced senescence by irradiating with 10Gy X-ray irradiation, or left the cells proliferating as controls. Next, either proliferating or irradiated GFP-positive cells were diluted at a ratio of 1:1000 into either proliferating or senescing cells that were negative for GFP (NEG) to achieve the following combinations: proliferating-GFP into proliferating-NEG; senescing-GFP into proliferating-NEG; senescing-GFP into senescing-NEG (Fig. 1A). We reasoned that in such a scenario, the proliferating-proliferating (Prol-Prol) conditions would resemble normal healthy conditions, the senescing into proliferating (Sen-Prol) would resemble low levels of senescence induction, while the senescing into senescing (Sen-Sen) would more closely resemble standard senescent culture conditions, or chronic senescent buildup as seen in damaged and aged tissue.

**Figure 1.**
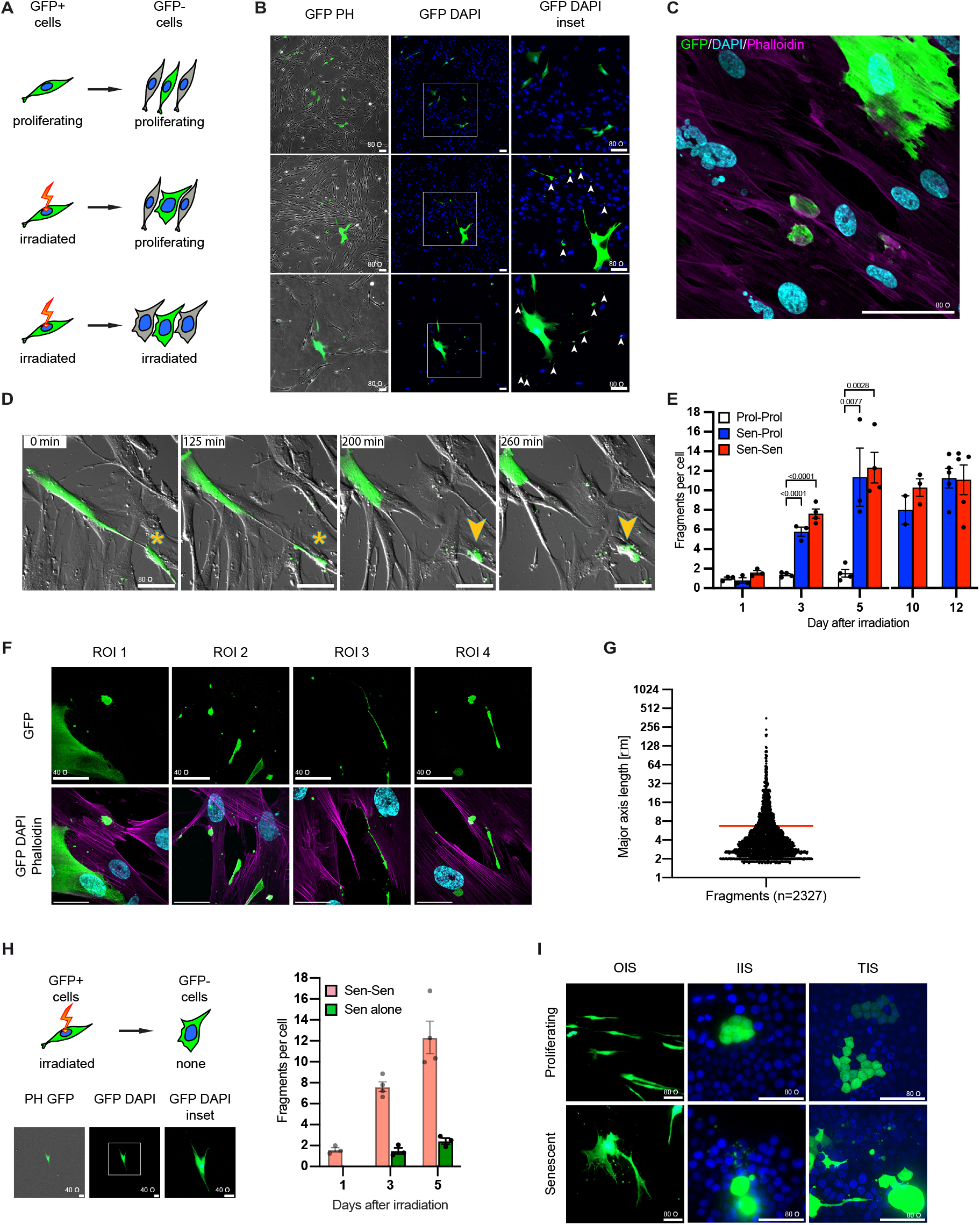
Cells undergoing senescence form cell-cell adhesion dependent fragments. (A) Experimental design of live cell imaging of individual GFP-labelled cells undergoing senescence. (B) GFP, phase contrast (PH) and DAPI images of cells diluted as per the scheme in A, on day 5 after plating. Arrowheads indicate GFP containing cellular fragments in the cultures with senescent cells (Scale bar = 80 µm) (C) Confocal image of senescent cell (green) diluted in GFP-negative cells, shows GFP fragments adhere to the membrane of adjacent cells in Sen-Sen condition. (Scale bar = 80 µm) (D) Timelapse imaging of GFP senescent cells diluted in GFP-negative proliferating cells at Day 3. Asterisk labels the adhesion point between cells. Arrowhead labels senescent fragment formation (Scale bar = 80 µm) (E) Quantification of fragments per cell at different time-points after senescence induction, in each culture condition. (Data are mean ± SEM, n = 3-5. One-way ANOVA, Dunnett’s multiple comparison test.) (F) Confocal micrographs (maximum Z-projection) of 4 different regions of interest (ROI) reveal the size and shape of senescent fragments. Top row shows anti-GFP immunostaining (green). Bottom row shows composites of chromatin (DAPI - cyan) and actin (phalloidin – magenta) staining and anti-GFP immunostaining (green). (Scale bar = 40 µm.) (G) Measurement of major axis length of fragments in cell-dilution cultures. (Red line indicates the mean of major axis length of all fragments, n=2327. Measurements were acquired from 3 biological replicates.) (H) Experimental design of live cell imaging of individual GFP-labelled senescent cells. GFP, phase contrast (PH) and DAPI images of individually plated cells (Scale bar = 40 µm). Graph shows quantification of number of fragments per cell (salmon bars show the same results as in E for reference, n = 3-4). (I) Images of cell-dilution strategy using oncogene induced senescence (OIS) in human primary lung fibroblasts (IMR-90) and irradiation-(IIS) and therapy-(TIS) induced senescence in human hepatocyte-derived carcinoma cell line (Huh7). (Scale bar = 80 µm, GFP, green; DAPI, blue)

Following irradiation and plating the various conditions, we imaged the cells each day. Unexpectedly, by 3-5 days post-irradiation, we repeatedly observed small GFP positive fragments in the cultures of cells induced to senesce, in both the proliferating and senescing environments (Fig. 1B). These were not visible, or were only present at very low levels in the proliferating control conditions. However, when we imaged these fragments at higher resolution, it was clear they were not pieces of cell debris in the media, but were GFP-positive cellular fragments bound to the membranes of adjacent cells (Fig. 1C). To determine the origin of these fragments, we next performed time-lapse imaging of the different culture conditions. In all conditions, cells routinely made contact with neighboring cells in a dynamic manner. In control cells (Prol-Prol), these contacts were non-permanent. Somewhat surprisingly, senescing cells also exhibited quite dynamic behavior, moving around the dish and making contacts. However, in cells induced to become senescent, sometimes these contacts persisted and the cells remained attached as the cells moved apart (Fig. 1D and Supplemental Movie 1). This movement resulted in the generation of long GFP-positive threads pulled between two cells, undergoing stretching and tension. Then, at one point, the senescent cells tore, leaving large GFP-positive fragments attached to the neighboring cell (Fig. 1D). Surprisingly however, following this process, both the fragments and the senescent cells remained intact, suggesting there are initial active membrane repair mechanisms present, even if the fragment may further disintegrate into smaller pieces over time. It appears that the strength of adhesion is quite high, as in some cases, there was clear recoil of the senescent cell following generation of the fragment. This process of adhesion-mediated fragmentation was consistently observed. We quantified the number of fragments at early timepoints, and found these became detectable around 3 days following irradiation, increasing to approx. 10-12 fragments per senescent cell at day 5 (Fig. 1E). This process of fragmentation continued, and similar numbers of fragments were seen at later stages in fully senescent cells (Fig. 1E). Together, this demonstrates that fragmentation is an ongoing and active process associated with senescent cells.

Next, we used image-analysis to quantify the fragment sizes. As these are dynamically-forming structures, there was great variation in the major-axis length, with some fragments stretching along the neighboring cell, while others had already rounded to a smaller size or disintegrated (Fig. 1F). Overall, omitting structures smaller than 2um, which may include secondary disintegration, this demonstrated that fragments are large structures with a mean length of the major axis of 7-8μm (Fig. 1G).

To investigate if cell-cell adhesion is required for this process, we next plated GFP-positive senescing cells at low density, in the absence of neighboring cells. These cells displayed similar senescent features as the previous conditions, but did not generate fragments (Fig. 1H). Together, this uncovers that senescent cells generate large cell fragments through a process of cell-cell adhesion. As a result, we chose to call these “senescent-cell adhesion fragments” (SCAFs).

As such cellular fragments had not previously been associated with senescent cells, we asked if they were unique to irradiation-induced senescence (IIS) in IMR-90 primary human fibroblasts. To address this we examined other conditions, including oncogene-induced senescence (OIS) in primary fibroblasts, or IIS or chemotherapy-induced senescence (TIS) in human liver cancer cells (Fig. 1I). In each case, SCAFs were detectable attached to neighboring cells. Similar fragments were also detected in mouse primary dermal fibroblasts (shown later in Extended data Fig. 6). Therefore, in every situation examined, senescent cells were found to undergo fragmentation.

### Senescent fragmentation is seen in vivo in OIS and aging endothelia

To further explore whether SCAFs are a defined structure associated with senescence, we wanted to examine if senescent cells in vivo exhibit features of fragmentation. To address this, we used a model of oncogene-induced senescence in mouse hepatocytes.^21^ Here, hydrodynamic tail-vein injection is used to deliver two plasmids, one with transposase and the other carrying a transposon with oncogenic NRas^G12V^-GFP or inactive NRas^G12V/D38A^-GFP. This results in construct expression specifically in individual hepatocytes in the liver ^21–23^ (Fig. 2A). In this setting, the hepatocytes expressing oncogenic Ras undergo senescence, while the inactive Ras-infected cells remain non-senescent, but in each case the infected cells are GFP-positive. The infected cells can then be identified with immunostaining for GFP, and additionally senescence markers such as SA-β-gal can be used to identify the senescent cells (Fig. 2A).

**Figure 2.**
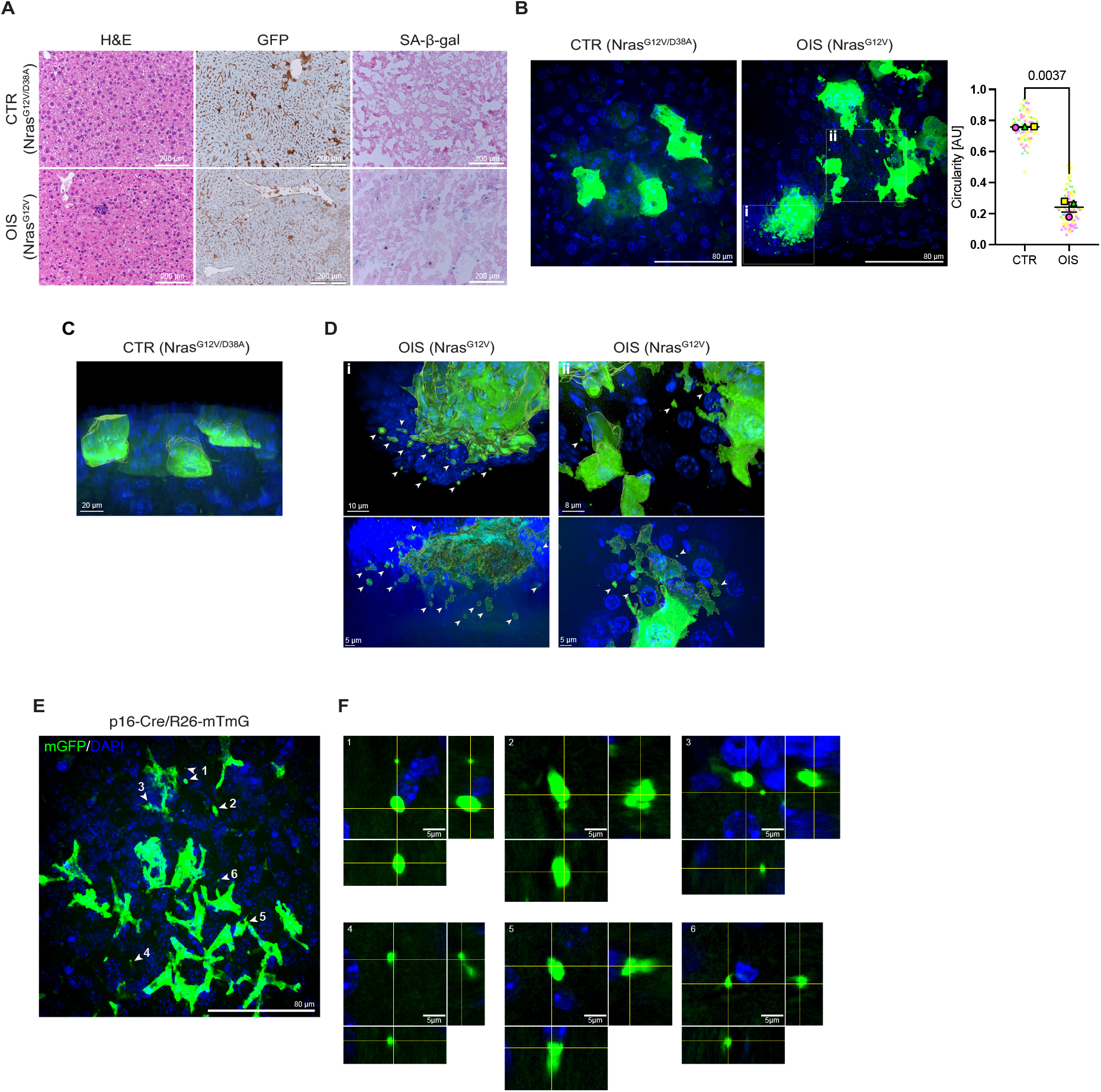
SCAFs are present in oncogene-induced senescent mouse hepatocytes and aging-induced senescent liver endothelial cells in vivo. (A) Hematoxylin Eosin (H&E), SA-B-gal staining and GFP immunostaining were performed on liver sections 6 days after hydrodynamic tail-vein mediated delivery of active Nras^G12V^ (OIS – oncogene induced senescence), or inactive Nras^G12V/D38A^ (CTR) and GFP coding sequences. (B) Maximum Z-projection confocal images of control (CTR, Nras^G12V/D38A^) and senescent (OIS, Nras^G12V^) hepatocytes expressing GFP. (GFP, green; DAPI, blue). Insets i and ii indicate regions of the image used for 3D rendering in the IMARIS software. (Scale bar = 80 µm) Graph shows quantification of cell circularity (Data are mean ± SEM, n = 3. Student’s t-test.) (C) IMARIS rendering of control hepatocytes (CTR, Nras^G12V/D38A^). (Scale bar = 20 µm) (D) IMARIS rendering of senescent hepatocytes (OIS, Nras^G12V^). White arrowheads indicate SCAFs not connected to senescent cell. (i scale bar = 10 µm, ii scale bar = 8 µm.) (E) Maximum Z-projection confocal image of senescent liver endothelial cells in one year old p16 reporter mice (p16-Cre/ROSA-mT/mG). Numbered arrows indicate SCAFs shown in F. (Scale bar = 80 µm) (F) Orthogonal views of SCAFs selected in E. (Scale bar = 5µm)

Next, on thick sections of liver containing the proliferating control or the senescent hepatocytes, we performed immunostaining and confocal optical-sectioning of GFP, generating image stacks (Fig. 2B). The control cells expressing inactive Ras-GFP clearly retained the epithelial shape of the hepatocytes, whereas the senescent cells displayed a much more irregular, less-circular shape, projecting many branches, as well as scattered non-connecting fragments (Fig. 2B). To determine whether these fragments were indeed separate from the senescent cells, or remained connected, we followed confocal imaging with 3D reconstructions. In each sample analyzed (n=3 livers), SCAFs were detectable adjacent to, but not connected to, the senescent cells, while none were obvious in the controls (Fig. 2C, D; Supplemental Movies 2-3). These results demonstrate that senescent cells *in vivo* generate SCAFs.

As a second approach to discern whether senescent cells undergo fragmentation in vivo, we investigated age-induced senescent cells, using a recently described senescence-reporter mouse model.^24^ Here, p16 positive cells are labeled with GFP, and have been demonstrated to increase in many aged tissues, including in endothelial cells in the liver. Through immunostaining and GFP imaging, we identified numerous GFP-positive senescent endothelial cells in the aged liver, which exhibited characteristics suggestive of cellular fragments (Fig. 2E). Employing confocal optical-sectioning, we were able to confirm that many of these were not physically connected to the senescent cells (Fig. 2F), supporting that senescent cells in aged tissues also undergo fragmentation.

### SCAFs contain a variety of organelles including damaged mitochondria

By imaging SCAFs in culture, we observed that these presented as large fragments of senescent cells that formed by attachment to neighboring cells. To further analyze the structure of these bodies, we performed correlative light and electron microscopy (CLEM). We plated senescing GFP-positive cells in an environment of GFP-negative senescing cells (Sen-Sen), and analyzed the cells at 5 days post irradiation. Fluorescence and brightfield imaging allowed us to identify and localize SCAFs of various sizes (Fig. 3A), and these samples were subsequently processed for EM analysis. The same SCAFs were readily identified at the ultrastructural level (Fig. 3B). In each case, the SCAFs appeared as large membrane-bound bodies, filled with cytoplasm. They contained a variety of organelles, including mitochondria, smooth endoplasmic-reticulum, and clathrin vesicles, as well as lipid droplets, autophagosomes and residual bodies (Fig. 3C). In most SCAFs, ribosomes were also clearly visible. In some instances, adherens junctions were visible at the site of adhesion between the SCAF and the neighboring cell, further supporting that SCAFs are generated by cell-cell adhesion (Fig. 3C, D). Quantification of the incidences of the main organelles validated their distribution patterns (Fig. 3E). Interestingly, in all analysis, we were unable to detect nuclei or nuclear material in any of the SCAFs analyzed.

**Figure 3.**
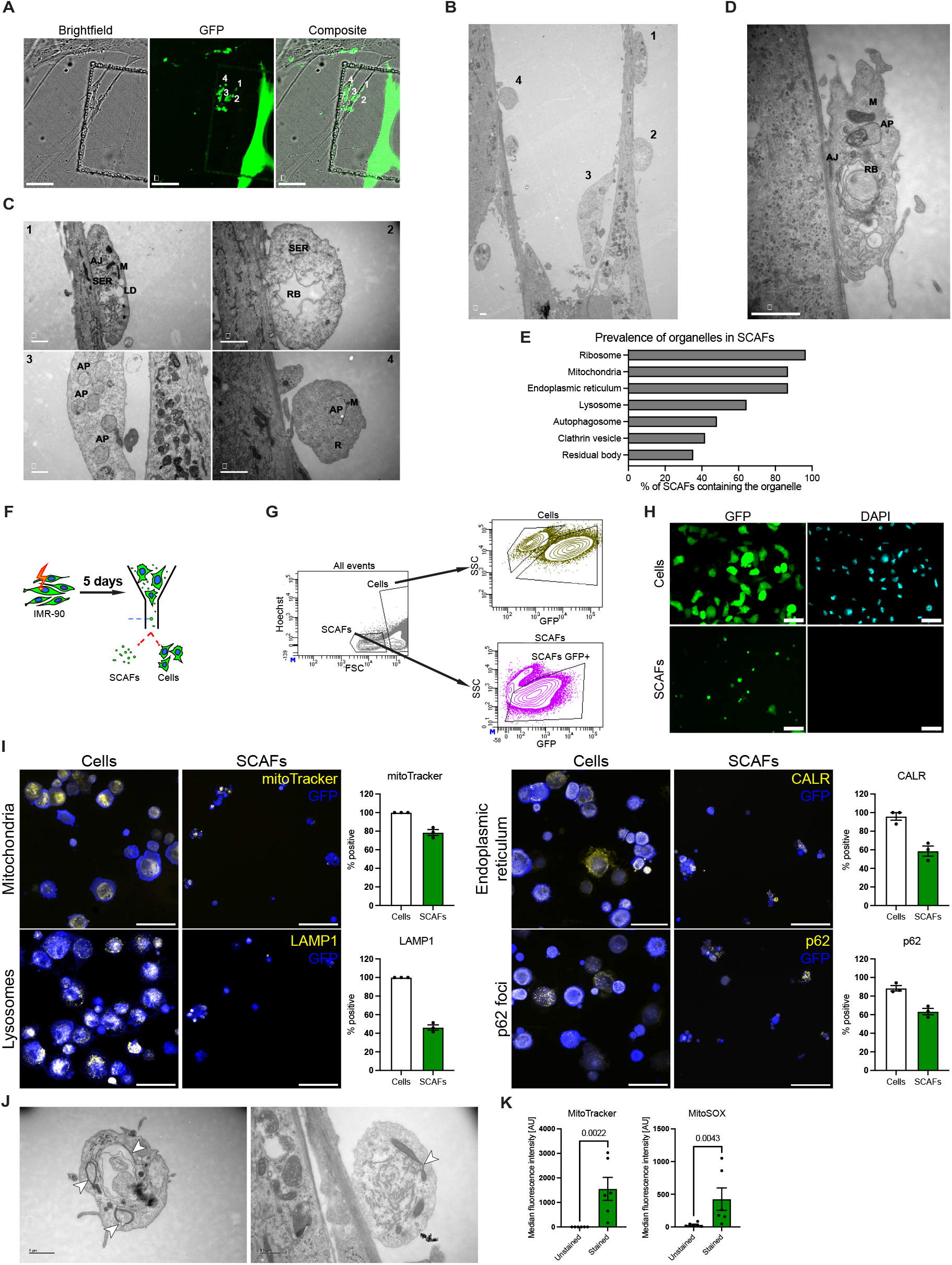
SCAFs are large membrane bound bodies containing damaged organelles. (A) Brightfield and GFP-image of live-cell microscopy of GFP-senescent cell dilution model. Numbers indicate individual SCAFs later imaged by TEM. (B) Low magnification TEM image of the region of interest shown in A. Four GFP positive SCAFs are indicated. (Scale bar = 1 µm.) (C) Higher magnification of GFP positive SCAFs 1-4 from B. (AJ, adherens junction: SER, smooth endoplasmic reticulum: M, mitochondria: LD, lipid droplet: RB, residual body: AP, autophagosome: R, ribosomes. Scale bar for 1,2 and 3 = 1 µm; for 4 is 0.5 µm.) (D) Additional SCAF showing adherens junction. (AJ, adherens junction: RB, residual body, AP, autophagosome: M, mitochondria. Scale bar = 1 µm.) (E) Percentage of SCAFs containing the indicated organelles (n=30). (F) and (G) Experimental design and gating of fluorescence-activated cell sorting of intact, GFP positive senescing cells and SCAFs (SCAF-FACS). (H) Cytospin and immunostaining of senescent cells and SCAFs for GFP (green) with DAPI staining of DNA (cyan). (Scale bar = 80 µm) (I) SCAF-FACS followed by cytospin combined with various staining approaches to visualize organelles present in senescent cells and SCAFs. Graphs indicate the proportion of cells/SCAFs containing the specified organelle. (n = 3, Scale bar = 80 µm) (J) Aberrant mitochondria are present in SCAFs as seen by CLEM. Arrows in left indicate a residual body and concentric laminated mitochondria with mitochondrial wall disruptions. Arrow in the right indicates a mitochondrial vacuole. (Scale bars = 1 µm and 0.5 µm respectively. (K) Graphs indicate the presence of mitochondria (MitoTracker) and mitochondria derived superoxide (MitoSOX) in FACS-analysed SCAFs. (n = 6, Mann-Whitney test)

To further investigate their structure and function, we devised a strategy to collect SCAFs through enzymatic dissociation and fluorescence-activated cell sorting (FACS) isolation (Fig. 3F, G). First, we isolated SCAFs and performed cytospin analysis, attaching the GFP-positive cells or SCAFs on glass slides (Fig. 3H). When stained for GFP and with DAPI to detect nuclei, we could not detect DAPI signal in the SCAFs, supporting our EM analysis that SCAFs do not contain nuclear material. Analysis of the size of these sorted SCAFs demonstrated they are similar to those formed in situ in the dilution assay (Extended data Fig. 2). Next, immunostaining for mitochondria, lysosomes and endoplasmic reticulum, again confirmed that in each case, these are frequently found in SCAFs (Fig. 3I).

Organelle, and in particular mitochondrial damage, has recently been highlighted as a prominent feature in senescent cells, contributing to the SASP.^25,26^ Interestingly, immunostaining for p62 foci, a marker of cellular damage and proteotoxic stress showed specific staining in senescent cells and SCAFs (Fig. 3I). Examination of the EM images of SCAFs also identified features of mitochondrial damage including evagination and blebbing (Fig. 3J). Measuring reactive oxygen species (ROS) activity, specifically mitochondrial superoxide, a marker of mitochondrial damage that is elevated in senescent cells, then confirmed the presence of damaged mitochondria (Fig. 3K). Altogether, this uncovers that SCAFs are large pieces of senescent cells, containing damaged organelles, in particular mitochondria, that become adhered to adjacent cells.

### Disruption of adherens junctions reduces SCAF formation, but increases cell death

A main question is whether SCAFs serve any functional role in senescent cells. To address this, we attempted to inhibit SCAF formation by disrupting adherens junctions. We used siRNA approaches to knockdown *CDH2* (N-cadherin) and *CTNND1* (p120 catenin) in senescent cells (Extended data Fig. 4). Then, GFP-positive senescent cells containing knockdown were diluted into GFP-negative proliferating cells. Quantification by either manual counting or FACS revealed a significant reduction in SCAF-incidence in both cases, demonstrating a functional role for cell-cell adhesion mediated by adherens junctions in the process of SCAF formation (Fig. 4B-D). Unexpectedly however, despite plating equal cell numbers after senescence induction and siRNA transfection, there were fewer cells remaining in the conditions with knockdown compared to siCTR (Fig. 4A). Measurement of cell-viability confirmed that loss of genes that facilitate SCAF formation, causes increased cell death in senescent cells (Fig. 4E), uncovering fragmentation as a protective mechanism in senescent cell survival.

**Figure 4.**
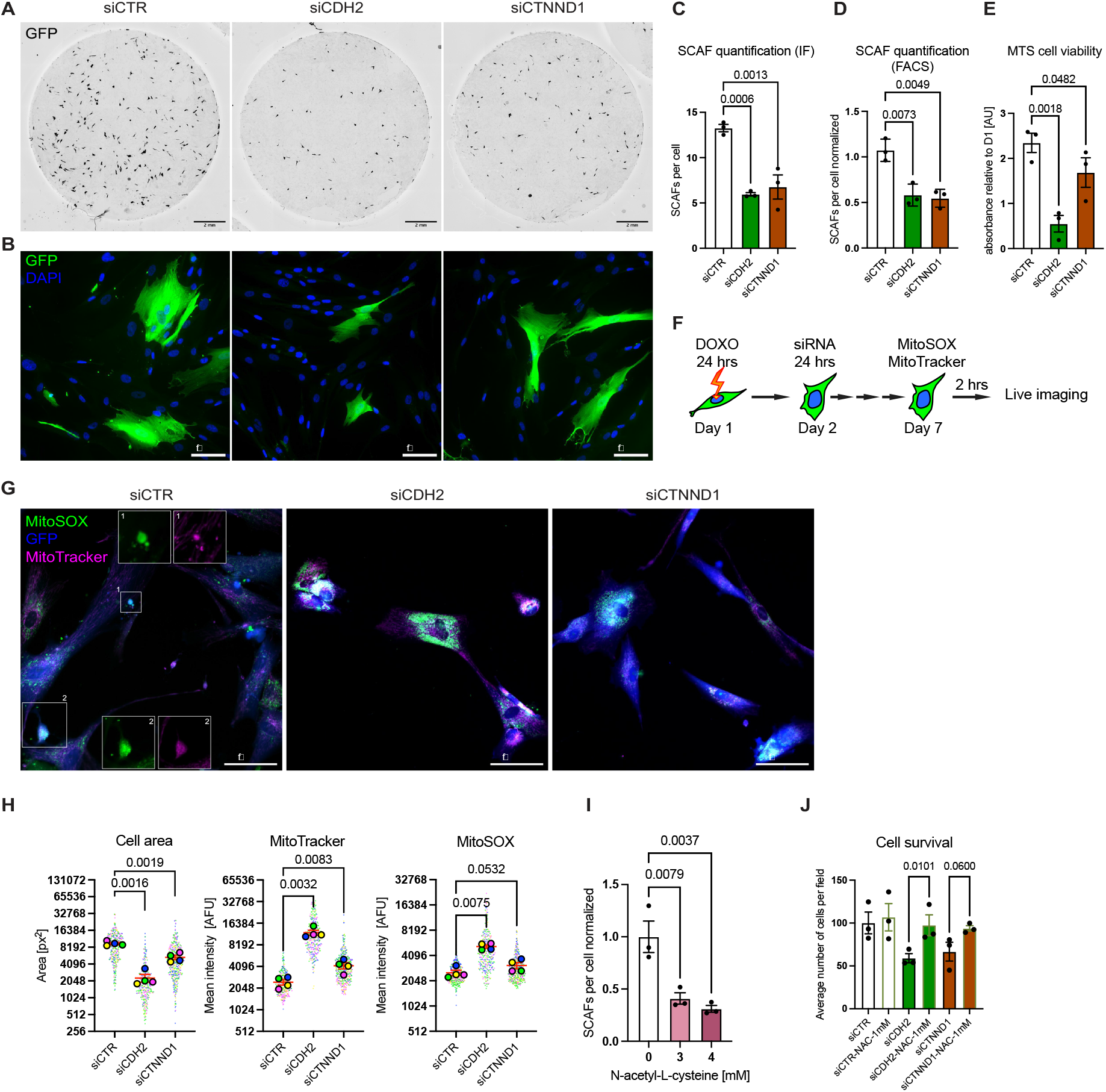
Knockdown of adherens junction (AJ) components in senescent cells attenuates SCAF formation and leads to reduced cell viability. (A) Whole-coverslip low-magnification scan of the GFP signal (black) in the cell-dilution model with adherens junction components knockdown: GFP+ IMR-90 cells induced to senesce (doxorubicin – DOXO), followed by siRNA treatment, were seeded with GFP-proliferating IMR-90 cells onto coverslips in equivalent numbers. Coverslips were stained and imaged 5 days after seeding. (Scale bars = 2mm.) (B) Higher magnification images from the samples shown in A (DNA – blue, GFP – green). siCDH2 (N-cadherin knockdown) and siCTNND1 (p120 knockdown) show reduced SCAF burden and smaller cell size after AJ components knockdown. (Scale bars = 80 µm.) (C) Quantification of SCAF formation by IF staining and manual counting on day 5 after seeding. (Data are mean ± SEM, n = 3. One-way ANOVA, Dunnett’s multiple comparison test.) (D) Quantification of SCAF formation by FACS analysis of the number of SCAFs and cells on day 5 after seeding. (Data are mean ± SEM, n = 3. One-way ANOVA, Dunnett’s multiple comparison test.) (E) Measurement of cell viability 4 days after siRNA transfection of senescent cells, using aqueous formazan-based colorimetric assay – MTS. (F) Experimental design of live cell imaging of mitochondria (mitoTracker) and mitochondrially produced superoxide (mitoSOX). Imaging is performed 4 days after siRNA transfection. (G) siCDH2 and siCTNND1 show striking differences in distribution and amount of mitoTracker (magenta) and mitoSOX (green) signal (GFP – blue) comp. Quantification in H. (Scale bars = 80 µm.) (H) Quantification of cell area and mean fluorescent intensities of mitoTracker and mitoSOX. (Data are mean ± SEM, n = 4. One-way ANOVA, Dunnett’s multiple comparison test.) (I) Quantification of SCAF formation by FACS analysis of the number of SCAFs and cells on day 5 after senescence induction. N-acetyl-L-cysteine treatment with indicated concentration was commenced right after induction and maintained until measurement. (Data are mean ± SEM, n = 3. One-way ANOVA, Dunnett’s multiple comparison test.) (J) Manual quantification of the number of cells per field of view (10 fields per condition) on day 5 after siRNA transfection. 1 mM N-acetyl-L-cysteine treatment rescues the cell loss caused by siCDH2 (and siCTNND1). (Data are mean ± SEM, n = 3. Two-way ANOVA, Šídák’s multiple comparison test – only indicated comparisons were performed.)

Mitochondrial damage must be regulated in senescent cells, as excessive damage can trigger cell death.^26^ Because we observed that SCAFs contain damaged mitochondria, we asked whether inhibiting SCAF formation would lead to the retention of damaged mitochondria within senescent cells. In control senescent cells, co-staining for mitochondria and ROS revealed detectable mitoSOX foci distributed throughout the cytoplasmic compartment, but with a clear enrichment in forming SCAFs (Fig. 4G). In contrast, cells treated with siCDH2 or siCTNND1 displayed visibly elevated mitochondrial ROS within the senescent cell body. Quantification confirmed these observations, with knockdown cells being smaller and accumulating higher levels of both mitochondria and ROS (Fig. 4H). These findings suggest that mitochondrial damage may act as a trigger for SCAF biogenesis.

To test this possibility, we sought to reduce mitochondrial damage by treating senescent cells with N-acetyl-L-cysteine (NAC), a ROS scavenger known to decrease mitochondrial stress in senescent cells. Strikingly, NAC treatment significantly reduced SCAF formation (Fig. 4I). Finally, to validate our model, we knocked down *CDH2* or *CTNND1* in NAC-treated senescent cells (Fig. 4J). Under conditions of reduced mitochondrial damage, loss of these proteins no longer induced cell death. Together, these results reveal that mitochondrial damage serves as an inducer of SCAF formation, and that SCAFs in turn promote senescent-cell survival by facilitating the removal of damaged organelles.

### SCAFs ultimately release their intracellular content, creating cellular debris

As the cell membranes of the SCAFs remain intact during their genesis, we wanted to explore the fate of these structures. To address this, we again sorted SCAFs, and now added them to cultures of proliferating IMR-90 fibroblasts, to track their behavior and dynamics (Extended data Fig. 5A). Interestingly, SCAFs remained intact throughout the procedure, and when plated in culture, they exhibited random dynamic behavior, which included spinning, projecting and retracting arms, or crawling-like behavior (Supplemental Movie 4). To determine how long the SCAFs persisted, we performed quantitative analysis and imaging over time, following their addition to cultures (Extended data Fig. 5B). During the first day, many SCAFs appeared intact, and tended to adhere to the cells in culture. After two days, there was a noticeable decrease in the number of SCAFs, but those remaining still clustered on cells. By three days, most of the GFP signal had disappeared, but any remaining signal was frequently attached to cells, in what appeared to be clusters of cellular debris (Extended data Fig. 5B). Imaging of individual SCAFs at these later timepoints showed that many of them were bursting, or undergoing lysis, and releasing their contents onto the attached cell and into the extracellular environment, while losing GFP-signal (Fig. 5A, Supplemental movies 5-6). To further examine if SCAF rupture generated debris, we repeated the sorting and addition of SCAFs to cultures, but this time adding higher numbers of SCAFs to the recipient cultures. As expected, after 3 days, this clearly resulted in large extra-cellular debris deposits (Fig. 5B).

**Figure 5.**
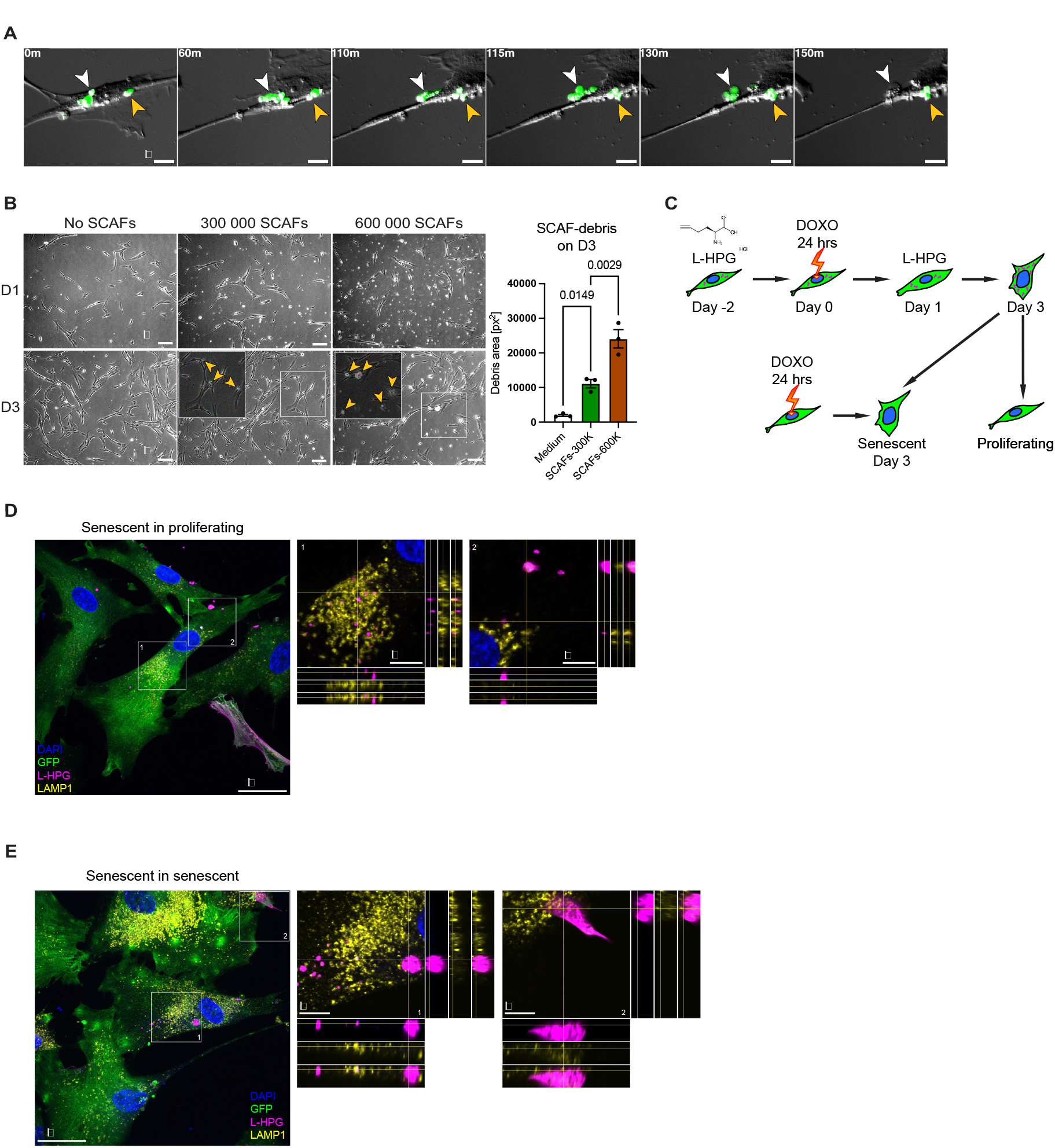
SCAFs maintain their integrity for a limited time after which they rupture, release their cytoplasmic content and form cellular debris. (A) Timeframe images of GFP positive (green) SCAF rupture and debris generation. Arrowheads indicate SCAFs at different time-points. (Scale bar = 10 µm) (B) Brightfield imaging at days (D) 1 and 3 after adding increasing number of SCAFs to culture generating increased cell debris (arrowheads). (Scale bar = 80 µm) Graph represents the quantification of the total debris area on day 3. (Data are mean ± SEM, n = 3. One-way ANOVA, Dunnett’s multiple comparison test.) (C) Experimental design of senescent cell nascent proteome labeling using L-homopropargyl glycine (L-HPG) shown in D and E. (D) and (E) Spinning-disc confocal imaging of L-HPG labelled GFP+ cells diluted in non-labelled GFP+. (DAPI – blue, GFP – green, L-HPG – magenta, LAMP1 – yellow.) In (D) labelled senescent are seeded with non-labelled proliferating cells. In (E) labelled senescent cells are seeded with non-labelled senescent cells. Orthogonal views of different channels are shown to demonstrate the distribution of the SCAF-derived material on the surface of, or inside of neighboring cells. (Scale bars = 80 µm)

This suggests that SCAFs can remain intact for a period of time, but then spill their contents onto the neighboring cells and into the extracellular space, suggesting a novel means of senescent cell communication. Even though GFP-signal was lost upon rupture, we speculated that the intracellular proteins might persist longer. To test this, we used a method to label the protein contents of the senescing cells and the SCAFs. We added L-Homopropargylglycine (L-HPG), a synthetic alternative to methionine that is incorporated during protein translation, and which can then be detected with click-chemistry (Fig. 5C). In this way, the protein content of senescing cells can be subsequently visualized.

We then diluted GFP-positive senescent cells treated with L-HPG, into GFP-positive L-HPG-negative cultures of proliferating or senescent IMR-90 fibroblasts. Then, these cells were stained using the click-reaction to visualize the L-HPG-labelled proteins, and immuno-stained for GFP and lysosomes (Fig. 5D, E). In each case, SCAFs staining positive for L-HPG were clearly evident, separated from the labeled senescent cells and attached to non-labelled ones. Interestingly, in both settings, intracellular L-HPG staining was evident, presenting as small regular dots, often co-localizing with LAMP1-positive lysosomes, suggesting that some protein from SCAFs was taken up and possibly degraded by the recipient cells (Fig. 5D, E). However, in most cases, the L-HPG labeled proteome from the SCAFs persisted as large extracellular accumulations, attached to the surface of neighboring cells. In many cases, such extracellular deposits were no longer GFP positive, indicating the SCAF had ruptured but its protein content remained as debris on the surface of the cell. This demonstrates that through SCAF formation and subsequent rupture, the intracellular content of senescent cells can be deposited onto, or taken up by, neighboring cells.

### The protein content of SCAFs includes DAMPs

The release of intracellular protein and organelles into the microenvironment can have diverse effects, depending on the content. To investigate the protein content of SCAFs that is presumably released upon rupture, we FACS isolated them at two different timepoints (days 5 and 10 after irradiation), and performed Mass Spectrometry (MS) analysis (Fig. 6A). This identified the SCAF-associated proteome, revealing strong overlap in protein content among the replicates, and identifying 1376 proteins common to the four replicates (Fig. 6B, Supplemental Table 1). Pathway analysis on these common proteins demonstrated signatures associated with immune cell recruitment and activation, in particular neutrophils, myeloid cells and T-cells (Fig. 6C). In addition, there was pronounced association with both NF-KB and Wnt signaling pathways. When we looked at KEGG pathway association, terms related to cell-cell adhesion appeared, including adherens junction, supporting our earlier findings, and reinforcing the mechanism of formation of SCAFs (Fig. 6D). Curiously, there was a strong association of the protein content with disease, in particular neurodegenerative disease, including Parkinson’s disease, Amyotrophic lateral sclerosis, Huntington’s and Alzheimer’s disease, all situations where senescence has previously been implicated. In order to determine if these signatures were conserved, we also performed similar MS proteomic profiling on SCAFs isolated from mouse dermal fibroblasts (Extended data Fig. 6). Interestingly, many of the same individual proteins and associated signatures seen in human fibroblasts, particularly those relating to neurodegeneration, were also detected.

**Figure 6.**
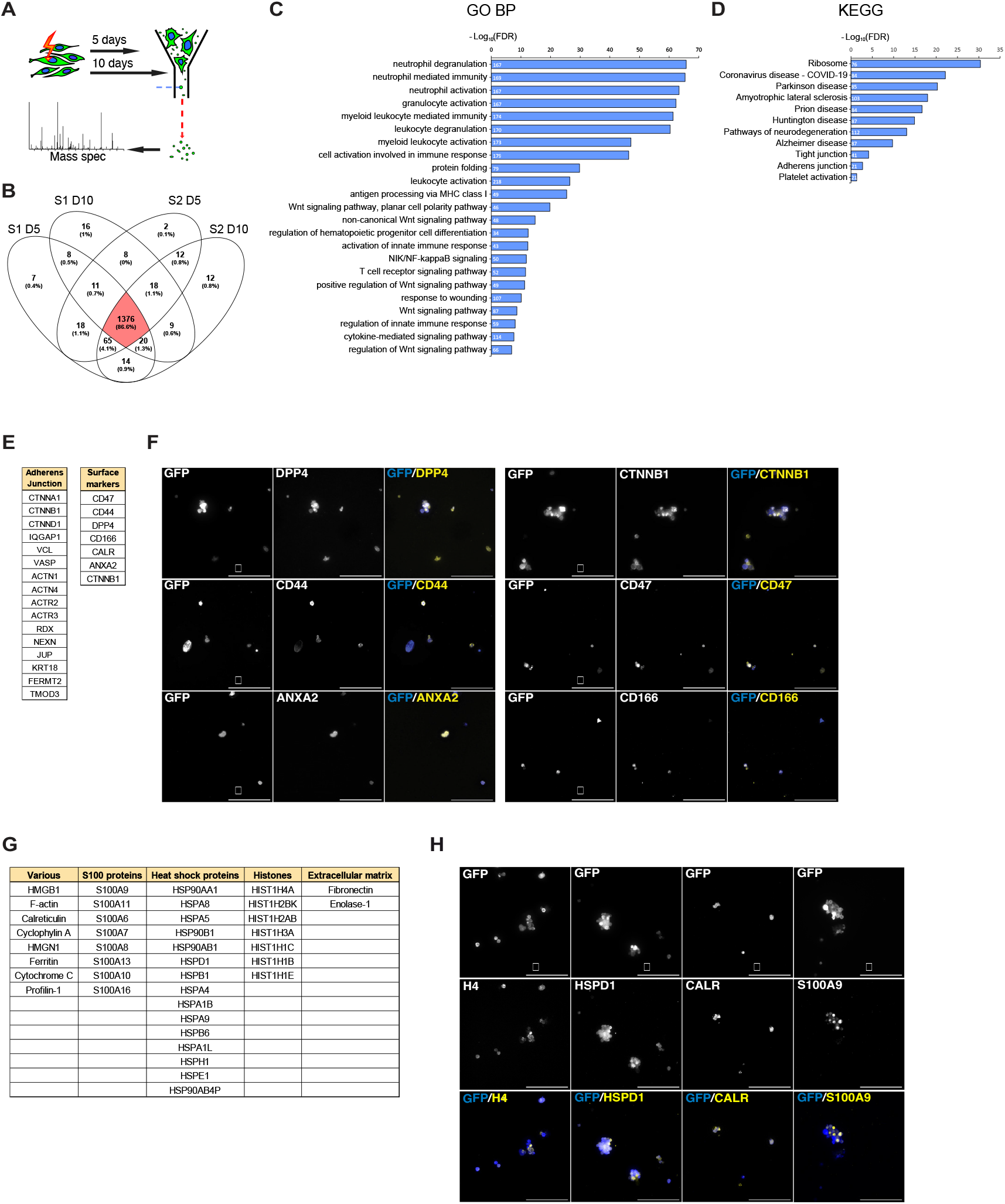
SCAFs contain proteins associated with innate immune processes, age-related diseases and contain Damage Associated Molecular Patterns (DAMPs). (A) Experimental design of Mass Spectrometry (MS) analysis of the protein content of SCAFs isolated from senescent IMR-90 cells. (B) Overlap of proteins identified in SCAFs by MS. (S1, sample 1; S2, sample 2; D5, day 5; D10, day 10). (C) Bioinformatic analysis of SCAF proteome using gProfiler (GO Biological process) reveals strong association with immune-related processes and signaling. (D) Bioinformatic analysis of SCAF proteome using gProfiler (KEGG pathway) reveals strong association with multiple age-related degenerative diseases. (E) Tables indicating protein components of adherens junction complexes and representative cell surface proteins detected in SCAF-proteome. (F) Immunofluorescence for representative surface marker proteins in isolated SCAFs. (Scale bar = 80 µm) (G) Table listing DAMP proteins detected in SCAF-proteome. (H) Immunofluorescence for selected DAMP proteins in isolated SCAFs. (Scale bar = 80 µm)

When we looked at individual proteins detected in the IMR90 cells, we observed most of the protein components of adherens junctions, including alpha- and beta-catenin, vinculin, VASP and others (Fig. 6E). In addition, there were many membrane proteins that have been associated with immune cell recruitment and senescence including calreticulin, CD44, CD47, and DPP4, a suggested marker of senescent cells ^27^ (Fig. 6E).We were able to validate our MS findings by immunostaining SCAFs from IMR-90 cells for many of these proteins (Fig. 6F).

It was also evident that many proteins known as damage-associated molecular patterns (DAMPs) are abundantly present in the SCAFs (Fig. 6G). These include calreticulin, ferritin, multiple Histones, heat shock proteins, cathepsins and high mobility group proteins, among others, findings which were also validated by immunostaining for individual examples on isolated SCAFs (Fig. 6H). This strongly supports that a complex mixture of organelles, as well as intracellular and membrane-associated proteins are released through SCAF formation and rupture.

### SCAFs activate cell migration and proliferation in normal and cancer cells

DAMPs play complex biological roles, influencing a variety of cell behaviors, including immune cell activation and recruitment, and cell migration and invasion linked to wound repair. We therefore sought to ask what effect SCAF exposure would have on cells. To address this, we performed RNA-sequencing on cells exposed to SCAFs. We isolated SCAFs from senescent cells and added them to proliferating IMR-90 cells for 20 hours, before sequencing the treated cells (Fig. 7A). Analyzing the RNA sequencing revealed that SCAF treatment resulted in 265 genes being significantly upregulated, and 316 genes downregulated (Fig. 7B). Pathway analysis on upregulated differentially expressed genes (DEGs) supported a strong correlation with SCAFs promoting proliferation, migration and invasion, with GO biological process terms such as *wound healing, regulation of cell motility, cell population proliferation*, p*ositive regulation of cell migration*, and *cell adhesion* associating with the upregulated genes (Fig. 7C). Analysis of KEGG associated terms revealed pathways associated with cancer, including *pathways in cancer, small cell lung cancer*, and *p53 signaling pathway*. Similar analysis on the downregulated genes further supported these changes, including GO biological process terms like *cell migration, cell motility, cell population proliferation* and *response to wounding* (Fig. 7D). Supporting this association, genes such as MMP1, MMP14 and MMP16 were induced by SCAFs, while genes for their potential substrates Collagens 1A1, 1A2, 3A1, 4A2, 5A1 and 5A2 were decreased.

**Figure 7.**
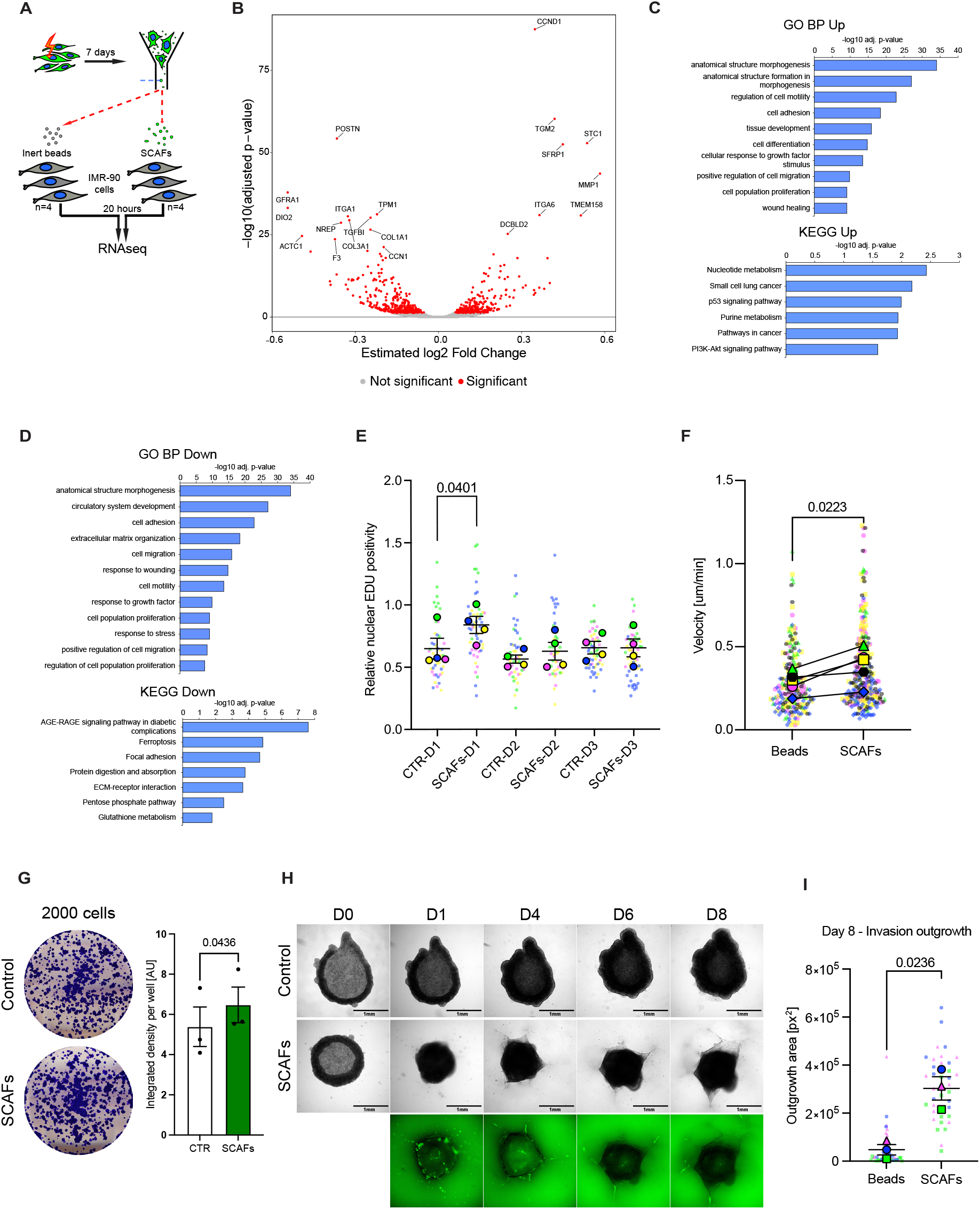
SCAFs promote proliferation, cell migration, and invasion of cancer cells. (A) Experimental design of RNA sequencing analysis of primary human fibroblast (IMR-90) after 20 hours of SCAF exposure. (B) Volcano plot of differentially expressed genes between IMR-90 cells treated with SCAFs compared to control cells treated with inert beads of similar size. (C) Bioinformatic analysis of significantly upregulated genes after SCAF-exposure using gProfiler. GO Biological process and KEGG pathway terms are shown. (D) Bioinformatic analysis of significantly downregulated genes after SCAF-exposure using gProfiler. GO Biological process and KEGG pathway terms are shown. (E) EDU incorporation assay at different time points (D, days) after SCAF-exposure in IMR90 fibroblasts. (Small dots indicate positivity of each field of view, colors represent biological replicates. Bordered dots indicate the mean of an experiment. Bars are mean ± SEM of proliferation relative to control cells on day 0. Two-way ANOVA, Šídák’s multiple comparison test – only indicated comparisons were performed.) (F) Cell motility measurement after SCAF-exposure in IMR-90 fibroblasts, relative to inert bead treated controls. (Small symbols represent median velocity of a single nucleus, colors indicate biological replicates. Bordered symbols show the mean of an experiment. Bars are mean ± SEM of the median velocity. Paired t-test.) (G) Focus formation assay of human hepatocyte-derived carcinoma cell line (Huh-7) on day 14 after treatment with SCAFs or medium on day 0 and day 2. Graph shows the quantification of the integrated density of the red channel of the whole well. (Data are mean ± SEM, n = 3, unpaired t-test.) (H) Huh-7 spheroid Matrigel-invasion assay after SCAFs or medium treatment. (Scale bar = 1 mm) (I) Quantification of Huh-7 invasion of Matrigel, measured as the outgrowth area of spheroids from H on day 8. (Small symbols represent outgrowth area of a single spheroid, colors indicate biological replicates. Bordered symbols show the mean of an experiment. Bars are mean ± SEM of the outgrowth area. Unpaired t-test.)

To determine whether functional changes were supported by these associations, we examined cells exposed to SCAFs. As expected from the sequencing results, exposure of IMR-90 fibroblasts to isolated SCAFs from senescent cells caused significant early increases in proliferation and migration (Fig. 7E, F).

As the SCAF proteome suggested significant communication with immune cells, we examined how SCAF-exposure would influence macrophages, which are frequently recruited to clear senescent cells.^21^ As we had performed for IMR-90 cells, THP1-derived M0 macrophages were exposed to SCAFs and then analyzed by bulk RNA sequencing (Extended data Fig. 7A). Here again, SCAFs activated similar genes and pathways in these cells, associated with migration and motility, but also differentiation, development and cancer (Extended data Fig. 7B-D). This further suggests that SCAFs may promote behavioral changes in cells consistent with wound repair.

As there was such a strong association with cancer signaling and invasion, and given that SCAFs are induced in senescent cancer cells, we also functionally investigated the effects of SCAF-exposure to cancer cells in culture. Here, we found that addition of SCAFs to liver cancer cells in culture also increased proliferation (Fig. 7G). Furthermore, when we added SCAFs to cancer cells in 3D culture, within days, the SCAF-treated cells invaded the gel (Fig. 7H, I). This supports our profiling and demonstrates that SCAFs can promote cell proliferation, migration and invasion in different contexts.

## DISCUSSION

Here we unveil a previously uncharacterized feature of senescent cells, cellular fragmentation. We have observed the detachment of sizable portions of senescent cells through cell-cell adhesion, coining the term “senescent cell adhesion fragments” (SCAFs). Critically, our investigation has revealed the presence of SCAFs across various senescent cell types in vivo and in vitro, suggesting that SCAFs may serve as additional markers of senescent cells. We identified the mechanism underlying SCAF formation, showing they arise from persistent adherens junction–mediated adhesion of senescent cells to neighboring cells, in the context of cell movement. Surprisingly, and contrary to the view of senescent cells as static, senescent cells in culture are highly dynamic, continuously moving and interacting with surrounding cells. Consistently, in vivo senescence often occurs in contexts of increased cellular dynamics, such as oncogene-induced hyperplasia or wound healing, where various cell types, even hepatocytes, can migrate,^28^ suggesting that SCAFs may contribute to these processes.

The discovery of SCAFs sparks questions into their nature and potential functions. Using various approaches, we identified that SCAFs contain a variety of organelles, including ribosomes, mitochondria and endoplasmic reticulum, and that the mitochondria exhibited typical features of damage previously described in senescent cells.^25,26^ However, despite this significant intracellular damage, senescent cells manage to survive, suggesting they have developed ways to tolerate this. This includes the release of mitochondrial DNA and dsRNA into the cytoplasm, preventing mitochondrial-damage associated cell death, but which contributes to SASP activation.^25,26^ We propose that that SCAF-mediated disposal of damaged mitochondria is another way of maintaining the level of mitochondrial damage in a tolerable range so as to maintain survival. These findings also introduce the concept that targeting SCAFs could offer a new strategy for eliminating senescent cells in senolytic approaches.

A major role of the SASP is to recruit immune cells, including macrophages, to clear the senescent cells. Our findings that SCAFs activate pathways of invasion and migration, also in macrophages, supports that this may be a complementary immune-recruitment property along with the SASP, but also initiated by mitochondrial damage. Interestingly, in senescent and damaged cells, damaged mitochondria are shuttled to the cell periphery, raising the possibility that such peripheral localization may favor their inclusion in SCAFs.^26,29^ The finding that inhibiting ROS and mitochondrial damage prevents SCAF formation supports such a cross-talk between mitochondrial stress and this adhesion-based fragmentation process.

Cells have developed various mechanisms to dispose of damaged mitochondria, including migrasomes and exophers. Migrasomes are small, organelle-containing vesicles deposited along the substratum as cells migrate.^29,30^ Exophers on the other hand are larger vesicles around 3.5um in diameter, also containing damaged mitochondria and protein aggregates but that are jettisoned to reduce toxicity, as shown in *C. elegans* neurons and mammalian heart.^31,32^ While sharing many apparent similarities with SCAFs, including a lack of nuclear material, these features are present in proliferating cells, and neither depends on cell–cell adhesion. SCAFs also appear larger than both. However, further studies directly comparing these structures and formation mechanisms may uncover further similarities or cross-talk, but altogether highlights the variety of complex ways cells have developed to dispose of damaged organelles.

In addition to uncovering SCAF-related mechanisms and functions, we also investigated the consequences of SCAF formation, uncovering deposition of a complex proteome including DAMPs. DAMPs, also known as alarmins or danger signals, encompass a diverse array of proteins, including heat shock proteins, S100 family members, annexins, histones, and calreticulin, as well as extracellular RNA and membrane lipids.^33–35^ When DAMPs become externalized, they promote proliferation, migration, and invasion, recruiting macrophages, neutrophils, and other immune cells, and play pivotal roles in wound repair.^34,36,37^ Our pathway analysis of the SCAF proteome and the transcriptional impact of SCAFs on cells aligns with these effects, highlighting the activation of signatures associated with wound repair and immune cell activation, and induction of proliferation, invasion, and migration in exposed cells. DAMPs are also central drivers of inflammaging, a chronic, sterile inflammation underlying many age-related diseases.^34,35,38^ As senescent cells are strongly associated with these conditions, the finding that senescent cells continuously dispose of DAMP-rich fragments suggests a direct mechanism by which they may actively fuel this persistent inflammatory state. Indeed, some DAMPs have been identified as SASP components, despite not being secreted via the known secretory pathways,^7,39^ suggesting that SCAF formation and rupture may be an additional mechanism by which these proteins are discharged and contribute to the SASP.

Finally, our study also reveals a robust association between SCAFs and cancer. SCAFs are induced in OIS and therapy-induced senescent cells, and the addition of isolated SCAFs to fibroblasts and cancer cells leads to activation of pathways and properties associated with tumor progression, including proliferation, migration and invasion. How the SCAFs promote these changes remains to be shown, but there are probably multiple factors involved. For example, CD44, present on SCAFs, can induce cancer cell proliferation through direct binding,^40^ while numerous DAMPs present in the SCAFs are also linked with cancer progression. We also show that SCAFs induce the expression of collagen-degrading enzymes. As matrix degradation is a primary feature of tumor invasion, there is a compelling rationale to further investigate whether the local production of SCAFs from age-associated or therapy-induced senescent cells could contribute to tumorigenesis and invasion.

Overall, we believe the identification of SCAFs uncovers yet another exciting feature in the complex biology of senescence, and the ways by which senescent cells interact with their microenvironment. As a core feature of senescent cells, which themselves become detrimental in chronic situations, heightened SCAF formation linked to an increased senescence burden is likely to have adverse effects associated with aging, cancer, neurodegeneration, and other diseases.

## MATERIALS AND METHODS

### Cells

All cells were cultured at 37°C in 5% CO2. IMR-90 cells, used at passages 11 to 15, were cultured in DMEM 41966 + 10% FBS + 100 U/ml penicillin-streptomycin. Huh-7 cells were cultured in RPMI 1640 w/10mM HEPES + 10% FCS + 2.5g/L Glucose + 1mM Sodium Pyruvate + 50 μM β-mercaptoethanol + 40 μg /mL gentamycine. All cell lines were routinely tested for the presence of mycoplasma.

### Mouse dermal fibroblast isolation

Primary cultured murine dermal fibroblasts (MDF) were obtained from C57BL/6, ROSA^mT/mG^ (mT/mG) and APP^NL-G-F/NL-G-F^ (APP) P0 pups. Animals were euthanized and the skin was placed in a sterile plastic dish (10 cm diameter), with the dermal side facing the dish, containing 2.5 mg/mL Dispase II (Roche) in PBS and incubated overnight at 4°C. Next day, the epidermis was separated from the dermis. The dermis was subjected to further digestion with 0.25% collagenase A (Roche) in PBS for 30 minutes at 37°C followed by 10 mg/ml DNase for 5 minutes. Dissociated cells were filtered through 70 μm cell strainer to generate a single-cell suspension and then collected by centrifugation and seeded onto 15 cm dishes and cultured in DMEM (4,5g/l glucose) w/GLutamax-I supplemented with 10% Foetal Calf Serum, AANE, 1mM Pyr Na 1 mM, 40 µg/mL gentamycine 100µM β-mercaptoethanol.

### THP-1 culture and differentiation into macrophages

THP-1 monocytes were cultured in suspension in RPMI 1640 w 10mM HEPES+ 10% FCS + Glucose 2,5 g/L (14 mM) + Sodium Pyruvate 1 mM + Gentamicine 40 µg/ml. To differentiate them into macrophages, cells were transferred into a culture dish and treated with 200 nM of phorbol 12-myristate 13-acetate for 24 hours, during which time they adhered to the bottom of the dish. After 24 hours, cells were washed in medium and cultured for another 24 hours in normal media. After this, cells were used depending on the experimental setup.

### Cell viral transduction

Transduction with retroviral vector was used to generate IMR-90 and Huh-7 cells stably expressing the green fluorescent protein (GFP). Retrovirus was produced by transiently transfecting the Phoenix packaging cell line (G. Nolan, Stanford University, Stanford, CA) with MSCV vector containing GFP coding sequence. Cells were infected with virus for 24 h, drug-selected in 1 µg/mL puromycin for 48 h, and subsequently tested for presence of replicatively competent virus. To generate IMR-90 cells stably expressing GFP and 4-hydroxy tamoxifen inducible ER:ras fusion protein, IMR-90 cells stably expressing GFP were transduced with a retroviral vector encoding the ER:ras protein (Narita, https://www.addgene.org/67844/). Cells were infected with virus for 24 h, selected for 7 days in 500 µg/mL G418 and subsequently tested for presence of replicatively competent virus.

### Induction of senescence

For ionizing radiation induced senescence, cells were exposed to a dose of 10 Gy of X-rays using CellRad Precision X-ray irradiator. The device was set to 130 kV and 5 mA at the tray position at one, leading to dose rate of 0.92 Gy/min.

For oncogene induced senescence, cells expressing the ER:ras protein (Young et al., 2009) were exposed to 500nM of 4-hydroxy tamoxifen for the full duration of the experiment. For therapy induced senescence, IMR-90 cells were treated with 250 nM doxorubicin for 24 hours and Huh-7 cells were treated with 200 nM doxorubicin for 48 hours. Afterwards, the culture was washed and cultured in regular medium. For dilution experiments, senescent cells were diluted in other cells as outlined. For quantification of SCAFs, the number of SCAFs in a given area was counted and expressed as a ratio of the number of senescent cells in the same area at the indicated times. This does not inform if each cell fragments at the same rate, but refers to total incidence at a given time.

### SA-ß-Galactosidase staining

IMR-90 stained for SA-ß-Galactosidase activity were washed and fixed in 0.5% glutaraldehyde (Sigma G7651) for 15 minutes at RT. Fixed cells were washed twice with PBS 1X containing 1 mM MgCl2 at pH6 and kept at 4°C until SA-ß-Gal experiment (no more than a week). Cells were stained in X-Gal solution consisting of 1 mg/mL X-Gal (Biosynth AG), 5 mM K3Fe(CN)6 and 5 mM K4Fe(CN)6 3H2O in PBS1X 1 mM MgCl2 at PH=6 for 8-12 hours. Cells were washed with PBS 1X and imaged with brightfield microscope (EVOS XL Core). SA-B-gal staining on liver tissue was performed on snap-frozen/cryosectioned tissues, that were fixed using glutaraldehyde and then counterstained with Eosin only.

### Knock-down using siRNA on Doxorubicin-treated IMR-90

siRNA pools of 5 siRNAs for human CDH2 (L-011605-00-0005), human CTNND1 (L-012572-00-0005) and non-targeting control siRNA (D-001810-01-05) were ordered from Dharmacon and were resuspended according to manufacturer’s instructions. Cells were seeded at 100,000 cells/well in 6-wells plate with 250 nM doxorubicin (Sigma, 44583-1MG) and incubated at 37°C and 5% CO2 for 24 hours. After doxorubicin removal, cells were washed with antibiotic-free medium containing serum. Transfection solution of siRNA and Lipofectamine RNAiMAX (Thermo Fisher 13778075) was prepared in serum-free and antibiotic-free Opti-Mem I and was incubated for 20 min at RT. Cells were then transfected with this solution in antibiotic-free medium for 24 hours, at final concentrations of 25 nM siRNA and 0.25% Lipofectamine. After 24 hours, cells were washed with regular medium and grown until desired day.

### Correlative light and electron microscopy (CLEM)

Fragmenting cells were cultured on laser micro-patterned Aclar supports (1,2). Using a spinning disc confocal microscope (Leica) a time-lapse acquisition was performed, followed by immediate fixation of cells and fragments using 2.5% glutaraldehyde and 2.5% formaldehyde in 0.1M cacodylate buffer for 1 hour at 4°C. After washing, the samples were post-fixed for 1 hour in 1% osmium tetroxide [OsO4] reduced by 0.4% potassium hexacyanoferrate (III) [K_3_Fe(CN)_6_] in H_2_O at +4°C. After extensive rinses in distilled water, samples were then stained by 1% uranyl acetate for 1 hour at +4°C, and rinsed in water. Samples were dehydrated with increasing concentrations of ethanol (25%, 50%, 70%, 90% and 3×100%), and embedded with a graded series of epoxy resin. Samples were finally polymerized at 60°C for 48 hours. Ultrathin serial sections (60nm) were picked up on 1% pioloform coated copper slot grids and observed with a Philips CM12 TEM electron microscope (Philips, FEI Electron Optics, Eindhoven, Netherlands) operated at 80kV equipped with an Orius 1000 CDD camera (Gatan, Pleasanton, USA).

### Fluorescence-Activated Cell Sorting (FACS) isolation

FACS was used to isolate senescent cells and SCAFs. Two million IMR-90 cells stably expressing GFP or mT/mG-MDF cells stably expressing tdTomato and seeded into a p15 petri dish were X-ray irradiated or doxorubicin treated and subsequently cultured for 5-10 days. Before sorting, cells were washed twice with PBS after which 4.5 ml of Accutase (STEMCELL technologies) was added, to dissociate cells and SCAFs and the dish was placed in an incubator. After five minutes cells were gently resuspended using a 5 mL serological pipette and incubated for 5 more minutes. Afterwards, 5.5 mL of 2% FBS in PBS was added and cells were again gently resuspended using 10 mL serological pipette. Then, cells were centrifuged at 290 rcf for 4 minutes. Supernatant was discarded and the pellet resuspended in 3 mL of Pre-Sort Buffer (BD biosciences). Gating for IMR-90 cells was set as in Fig. 4A and similar strategy gating for tdTomato was used for mTmG-MDF cells. Fragments or cells were sorted at RT and collected into 1.5 mL Eppendorf tubes with 350 µl of standard media.

### Flow-cytometry analysis

For flow-cytometric quantification of SCAFs per cell (FACSquant), cells were first pretreated with Hoechst 33342 in the dish by adding a stock solution to achieve final concentration of 800 nM. The dish was returned to the incubator for 20-minute incubation. Afterwards, cells were processed the same way as for FACS isolation but resuspended in 0.5ml PS buffer in the last step.

Flow-cytometry analysis was performed on BD Symphony under FSC threshold settings of 500. All events were first gated in FSCxSSC to large (cells) and small (debris). Cells were then gated for GFP and Hoechst positivity and GFP+ cells were counted. Debris was then gated for GFP positivity and Hoechst negativity and SCAFs were counted. The gate for GFP positivity was determined beforehand by comparison with GFP-senescent IMR-90 cells. Around 5000 cells were recorded for each analysis.

Flow cytometric analysis for the presence of mitochondria and mitochondrial superoxide radicals in SCAFs was performed in a similar manner as FACSquant. First cells were washed with HBSS (w Ca, Mg), pretreated with mitoSOX Red (1μM) and mitoTracker Deep Red FM (50nM) in HBSS in the incubator for 30 minutes. Then cells were processed and analysed similarly as in FACSquant. The GFP+ SCAFs were then analyzed for mitoSOX (PE-TexasRed) and mitoTracker (APC) signal median fluorescence intensity.

### Cytospin

FACS sorted cells or SCAFs were fixed in 4% PFA for 15 minutes at RT and then washed 3x by PBS. Suspension of 150 µL was added into a cytospin funnel mounted on poly-lysine coated microscope slide and centrifuged at 1000 RPM for 5 minutes. Afterwards, the sample was fixed in 4% PFA for 15 minutes at RT, to attach the sample to the slide surface.

### Immunofluorescence

Cells/fragments for immunofluorescence were cytospun (see above) or fixed in 4% PFA for 15 minutes at RT followed by 3x PBS wash. Next, the sample was permeabilized for 10 min in PBS + 0.1% Triton-X and washed 3x by PBS. Blocking was done using PBS + 10% goat serum + 0.1% Tween for 1 hour at RT. Primary and secondary antibodies were diluted in PBS +1% goat serum + 0.1% Tween. After blocking, sample was incubated with primary antibody for 1 hour at RT, followed by 3x PBS wash. Then, secondary antibody was added for 1h at RT followed by 3x PBS wash. . Specifically in the case of staining with anti-LAMP1 antibody, the permeabilization, blocking and staining was done with 0.05% saponin (Thermo Scientific, J63209.AK) and without Triton-X or Tween. Subsequently, nuclei were counterstained using DAPI 1 µg/ml for 10 minutes at RT, and actin filaments were counterstained using Phalloidin-AF568 (ThermoFischer) 165 nM. Sample was then washed 3x by PBS and mounted using Fluoromount-G (ThermoFischer).

For immunofluorescent imaging, livers were fixed with 10% neutral buffered formalin for 24 hrs at 4°C and stored in 70% EtOH until further processing. Fixed liver tissues were washed in PBS and sectioned at 100 µm using vibratome. Sections were permeabilized using PBS + 0.5% Triton-X at 4°C overnight. Blocking was done at 4°C for 24 hours in PBS + 10% goat serum + 0.1% Triton-X and primary and secondary antibodies were dissolved in PBS+1% goat serum +0.01% Triton-X. Following blocking, sections were incubated in primary antibody for 24 hours at 4°C, followed by 3x 2-hour wash at 4°C in PBS + 0.01% Triton-X and 24-hour incubation at 4°C with secondary antibody. Afterwards, sections were stained with DAPI 1 µg/ml for 1 hour at 4°C and washed in PBS + 0.01% Triton-X for 3x 2-hours at 4°C. Sections were then mounted on a microscope slide and cleared by serial treatment with 60%, 80% and 87% glycerol for 2 hours per step. Afterwards, samples were mounted in 87% glycerol.

Primary antibodies were used at following concentrations: anti-GFP (1:1000, Abcam, ab13970), anti-DPP4 (1:300, Abcam, ab28340), anti-CTNNB1 (1:500, Abcam, ab16051), anti-CD44 (1:1000, Abcam, ab15710), anti-CD47 (1:1000, BioXcell, MIAP410), anti-ANXA2 (1:500, BD biosciences, 610069), anti-CD166 (1:500, Abcam, ab109215), anti-Histone H4 (1:300, Abcam, ab10158), anti-HSPD1 (1:300, In-House produced, clone: 4MTE-2H7), anti-CALR (1:500, Abcam, ab2907), anti-S100A9 (1:1000, Abcam, ab92507), anti-p62 (1:1000, Abcam, ab109012), anti-LAMP1 (1:300, Abcam, ab24170). Alexa-conjugated secondary antibodies were used in the dilution of 1:1000 – 1:2000.

For mitoSOX Red (1μM; Invitrogen M36008) and mitoTracker Deep Red FM (50nM, Invitrogen M22426), cells were washed with HBSS (w Ca, Mg) and pretreated in the same for 30 minutes at 37°C, 5% CO_2_ (incubator). Then cells were washed 3x with HBSS (w Ca, Mg) and imaged in the same medium. Cells were imaged at 37°C, 5% CO_2_. MitoSOX Red was imaged using the excitation wavelength of 385nm and the emission was collected with a dsRed filter (592/20 nm).

### L-HPG whole proteome labeling

Proliferating GFP labelled IMR-90 cells were treated with 200 µM L-homopropargylglycine (L-HPG) in DMEM without Methionine and cystine medium for 48 hrs. Afterwards, cells were split and reseeded with 250 nm doxorubicin for 24 hours to induce senescence. Then cells were again treated with 200 µM L-HPG in methionine free medium for 48 hrs. Afterwards, cells were trypsinised and diluted/seeded (1:1000) with proliferating or senescent GFP+ IMR-90 cells in glass-bottom dishes. The cells were fixed 5 days after seeding. The click-it staining protocol is the same as for EDU proliferation assay followed by GFP immunostaining (see below and above respectively).

### EDU proliferation assay

Cells were incubated with 20 µM of EDU in standard medium for 2 hours. Subsequently, cells were 1x washed with PBS and fixed for 15 minutes in 4% PFA at RT. After fixation, samples were incubated with 100 mM Tris (pH 7.5) for 5 minutes and permeabilized with PBS + 0.1% Triton-X for 10 minutes. Following 3x wash with PBS, cells were stained using Click-staining solution containing either Sulfo-Cy3-Azide or Alexa-Fluor-555-azide for 30 minutes at RT. Subsequently, samples were washed 3x with PBS, stained with DAPI 1 µg/ml for 10 minutes at RT, washed 3x with PBS and mounted using Fluoromount.

### MTS assay

Cell viability was measured using CellTiter 96® AQueous Non-Radioactive Cell Proliferation Assay (MTS) (Promega, G5421) according to the manufacturer’s instructions.

### Time-lapse and confocal microscopy

For acquisition of in-vitro senescent cells and senescent hepatocytes in liver sections, Leica CSU-W1 spinning disk, 63× objective, oil immersion was used with Z step of 0.3 µm. For confocal images of fragmenting cells Leica SP5 inverted confocal microscope was used with 63x objective, oil immersion.

For live-cell imaging cells were seeded in a 30 mm diameter glass-bottom dish. For acquisition, Zeiss Axio Observer Z1 microscope with 20x dry objective was used. Cells were treated with 30 000 SCAFs and were placed into a humid heated chamber with 5% CO2 and 80% humidity in the microscope. Four positions were selected for each condition and pictures were taken every 5 minutes for 8 hours.

### Mass spectrometry analysis of SCAF proteome

After FACS isolation, GFP positive SCAFs from senescent IMR-90 cells were centrifuged at 400g for 5 minutes at room temperature and the pellet was snap-frozen in liquid nitrogen. Same approach with adjusted sorting strategy was used for tdTomato positive SCAFs from senescent MDF cells isolated from mTmG mouse line. Proteins were then precipitated using trichloroacetic acid, followed by acetone wash. Mass spectrometry was performed on Orbitrap Elite using the Top20CIDr method, with 2 hours run and 3 repetitions. Results were analyzed using Proteome Discoverer 1.4 and compared with homo sapiens_181015 database. Thresholds of 1% FDR and minimum of 2 peptides per protein were applied. Pathway analysis was performed using gProfiler (https://biit.cs.ut.ee/gprofiler/gost). Gene Ontology Databases and KEGG biological pathway database were considered. Adjusted p-value of <0.05 was used as a threshold to select the significant enrichment.

### RNA sequencing of SCAF treated cells

Proliferating GFP negative IMR-90 cells were treated with 300 000 SCAFs or inert beads for 20 hours under normal culture conditions. Four independent biological replicates per condition were generated. The same approach was used for THP-1 derived macrophages, which were done in three biological replicates per condition. Cells were washed 1x with PBS and lysed in 350 µl of Guanidine thiocyanate-based lysis buffer (LBP lysis buffer, Macherey-Nagel NucleoSpin RNA plus isolation kit, Ref. 740984), RNA was isolated and sent to the GenomEast platform at Institut de Génétique et de Biologie Moléculaire et Cellulaire (IGBMC). Library was prepared using TruSeq Stranded mRNA Library preparation (Illumina). Sequencing was performed with HiSeq 4000 technology (Illumina) on single-end 50bp length reads. Reads were pre-processed to remove low quality and short (<40bp) reads using cutadapt v1.10. Remaining reads were mapped onto the hg38 assembly of Homo sapiens genome using STAR version 2.5.3a. Gene expression quantification was performed from uniquely aligned reads using htseq-count version 0.6.1p1, with annotations from Ensembl version 98 and “union” mode. Read counts have been normalized across samples with the median-of-ratios method proposed by Anders and Huber, to make these counts comparable between samples. Differential gene expression analysis was performed using Wald test, as proposed by Love et al. and implemented in the Bioconductor package DESeq2 version 1.16.1. P-values were adjusted for multiple testing using the Benjamini and Hochberg method. Genes with an adjusted p-value <0.05 were considered significantly differentially expressed. Pathway analysis was performed using gProfiler (https://biit.cs.ut.ee/gprofiler/gost). Gene Ontology Databases and KEGG biological pathway database were considered. Adjusted p-value of <0.05 was used as a threshold to select the significant enrichment.

### Cell motility assay

Proliferating IMR-90 cells were seeded in a 4 chamber 30 mm diameter glass-bottom dish (IBL, Ref 220.120.022) with 8000 cells per chamber. For acquisition, Zeiss Axio Observer Z1 microscope with 20x dry objective was used. Cells were treated with Hoechst 33342 0.5 µg/ml, 30 000 SCAFs or inert beads and were placed into a humid heated chamber with 5% CO2 and 80% humidity in the microscope. Four positions were selected for each condition and pictures were taken for 8 hours every 5 minutes. Cell nuclei were automatically detected using StarDist and subsequently tracked using the TrackMate ImageJ modules.

### Focus formation assay

Huh-7 cells were seeded into 6 well plates in two different densities – 2000 cells and 4000 cells per well. At 24 and 48 hours after seeding, cells were treated either by normal medium or by 60 000 SCAFs isolated from senescent IMR-90 cells expressing GFP. After 10 days, cells were fixed in 4% PFA at RT for 15 minutes and washed with PBS. Subsequently, cells were stained with crystal violet using a standard protocol.

### Spheroid Matrigel-invasion assay

To generate spheroids derived from Huh-7 cells, 40 000 cells were seeded into Ultra-low adhesion 96 well plates (Corning). After 48 hours, formed spheroids were treated either with standard medium or 30 000 SCAFs. Plate was then centrifuged at 300g for 5 minutes. Supernatant was aspirated and spheroids were embedded in Matrigel +10% rat tail collagen. Spheroids were imaged daily using a 5x dry objective. Eight technical replicates were performed in each independent experiment. On day 8, the area of visible outgrowth was manually traced and measured.

### Statistical analysis

Unless otherwise specified, results from each group were averaged and expressed as means + standard error of the mean (SEM) (represented as error bars). T-test was used for comparison between 2 groups and one-way ANOVA analysis was used for comparisons among 3 or more groups. Two-way ANOVA analysis was used to examine the influence of two different independent variables. The Statistics program Prism (GraphPad Prism 9) was used for analysis.

## ACKNOWLEDGEMENTS

This work was supported by grants from La Fondation pour la Recherche Medicale (FRM) Amorcage pour les jeunes equipes (AJE20160635985), and Programme “Maladies Neurodégénératives 2020” (MND202004011743) ; Fondation ARC pour la Recherche sur le Cancer (PJA20181208104) ; IDEX Attractivite′ - University of Strasbourg (IDEX2017) ; La Fondation Schlumberger pour l’Education et la Recherche FSER 19 (Year 2018)/FRM ; Agence Nationale de la Recherche (ANR) ANR-19-CE13-0023-03 and ANR-22-CE14-0062-01 ; la Fondation pour la Recherche sur Alzheimer (2024) ; and Ligue Contre le Cancer (all to W.M. K.). The work was also supported by an institutional grant to the IGBMC, ANR-10-LABX-0030-INRT, a French State fund managed by the Agence Nationale de la Recherche under the frame program Investissements d’Avenir ANR-10-IDEX-0002-02. Sequencing was performed by the GenomEast platform, a member of the “France Genomique” consortium (ANR-10-INBS-0009). We acknowledge the support of the Light Microscopy Facility at the IGBMC imaging center, member of the national infrastructure France-BioImaging supported by the French National Research Agency (ANR-10-INBS-04). The funders had no role in study design, data collection and analysis, decision to publish, or preparation of the manuscript. We thank Katerina Jerabkova-Roda for valuable feedback on the work and sharing resources.

## FIGURE LEGENDS

**Supplementary Figure 1.**
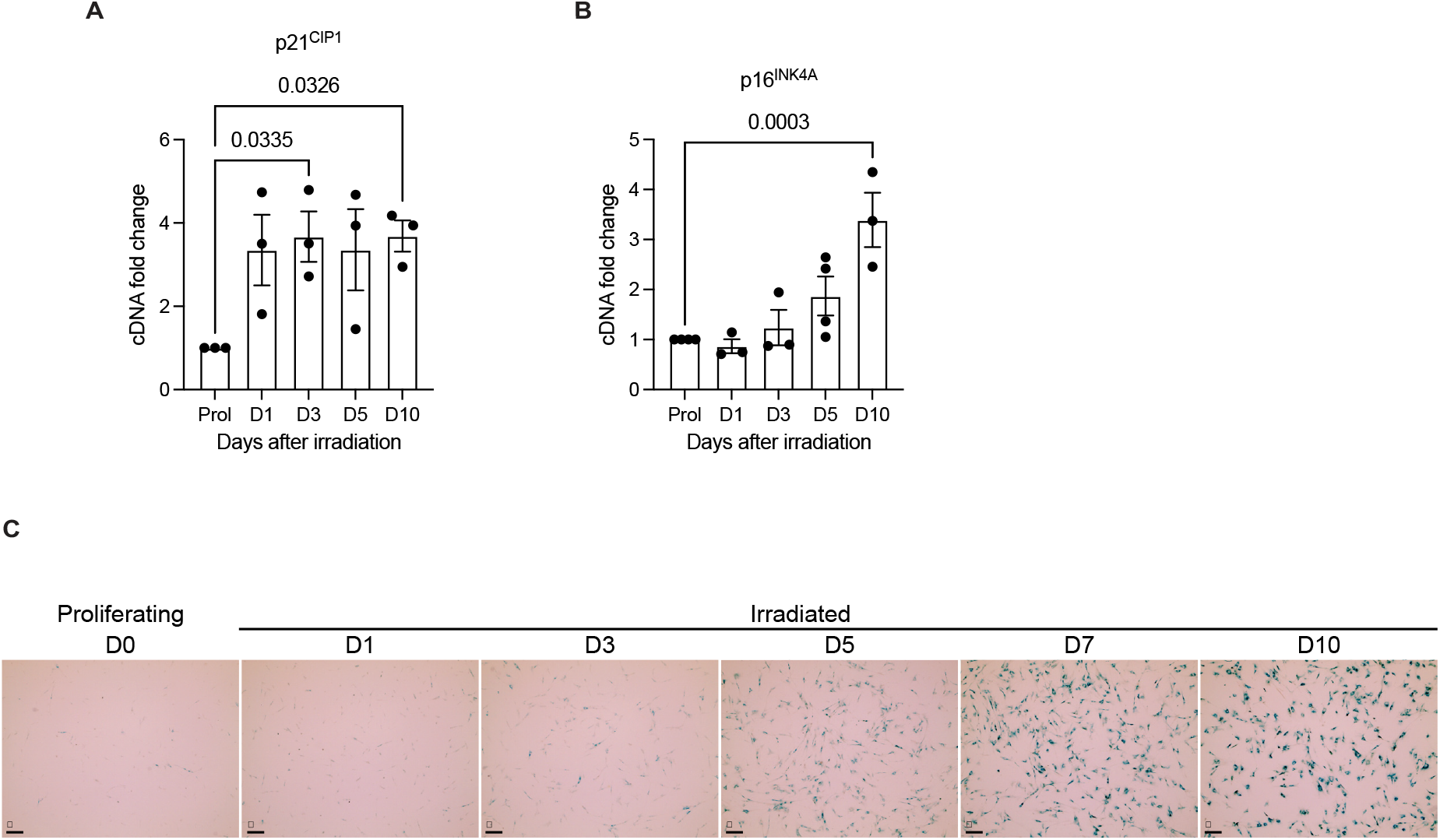
The dynamics of senescence features in irradiation-induced senescence. (A) Time-course of the expression of p21^CIP1^ after senescence induction in IMR90 cells. (Data are mean ± SEM, n = 3. One-way ANOVA, Dunnett’s multiple comparison test.) (B) Time-course of the expression of p16^INK4A^ after senescence induction in IMR90 cells. (Data are mean ± SEM, n = 3-4. One-way ANOVA, Dunnett’s multiple comparison test.) (C) Time-course of SA-B-gal positivity after senescence induction in IMR90 cells.

**Supplementary Figure 2.**
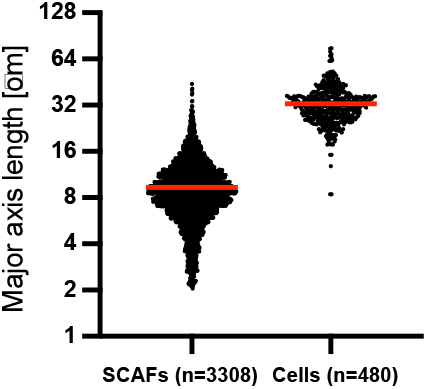
Average major-axis length of FACS-isolated senescent cells and SCAFs.

**Supplementary Figure 4.**
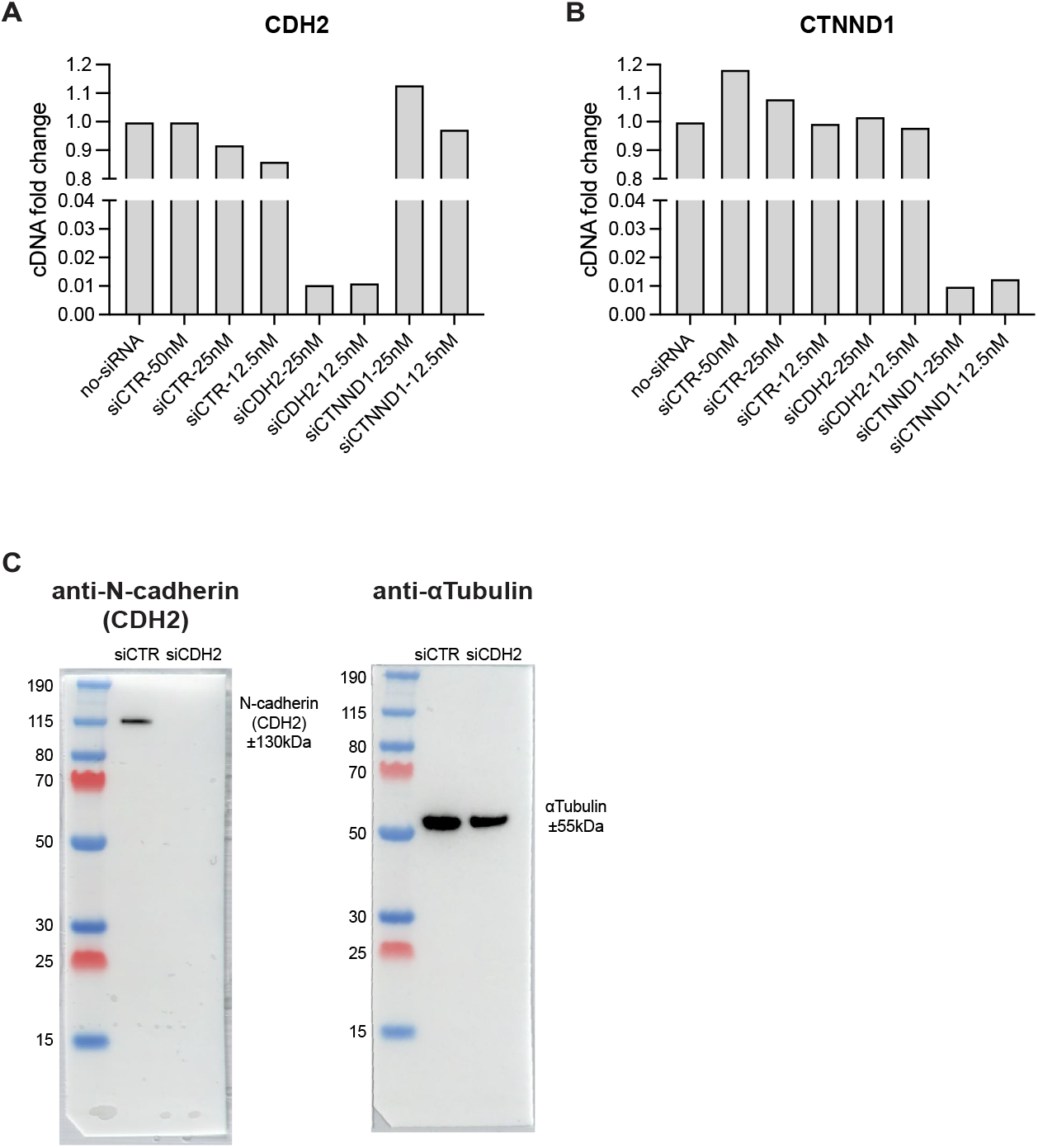
Validation of siRNA approaches targeting adherens junction components. (A) Relative mRNA levels of CDH2 gene targeted by the corresponding siRNA. Normalized to Rplp0 and no-siRNA. n=1 (B) Relative mRNA levels of CTNND1 gene targeted by the corresponding siRNA. Normalized to Rplp0 and no-siRNA. n=1 (C) Western blot for N-cadherin (encoded by the CDH2 gene) and Tubulin after targeting with 25 nM siRNA. n=1

**Supplementary Figure 5.**
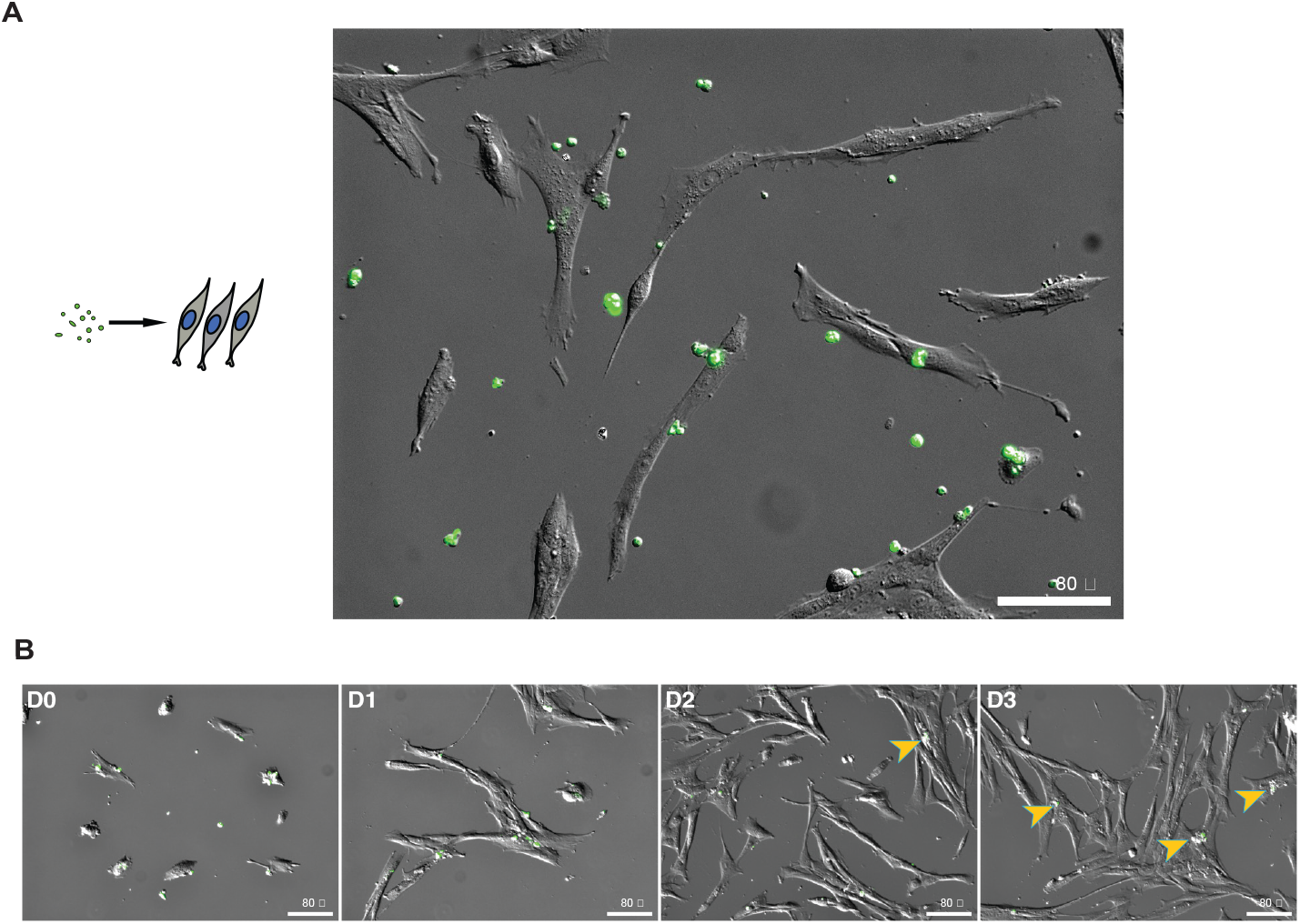
FACS isolated SCAFs maintain their integrity for several days. (A) Image of living GFP positive (green) SCAFs isolated by FACS and seeded on top of proliferating GPF-negative IMR-90 cells. (Scale bar = 80 µm) (B) Representative images of GFP positive (green) SCAFs at different times (Days0-3) after seeding on proliferating GPF negative IMR-90 cells. Arrowheads indicate residual GFP positive SCAFs surrounded by GFP negative cellular debris. (Scale bar = 80 µm)

**Supplementary Figure 6.**
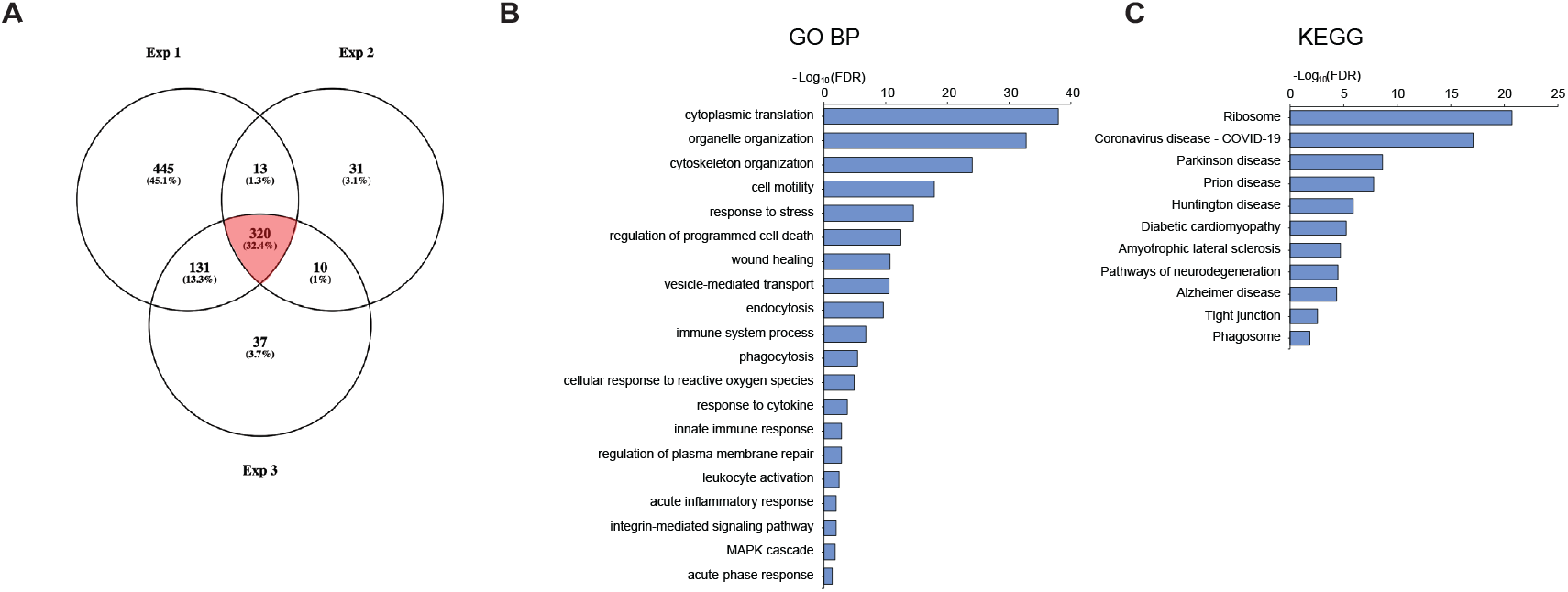
Mass Spectrometry analysis of the protein content of SCAFs isolated from senescent mouse dermal fibroblast (MDF) cells. (A) Overlap of proteins identified in MDF derived SCAFs by MS. (B) Bioinformatic analysis of MDF SCAF proteome using gProfiler (GO Biological process) reveals strong association with immune-related processes and signaling. (C) Bioinformatic analysis of MDF SCAF proteome using gProfiler (KEGG pathway) reveals strong association with multiple age-related degenerative diseases.

**Supplementary Figure 7.**
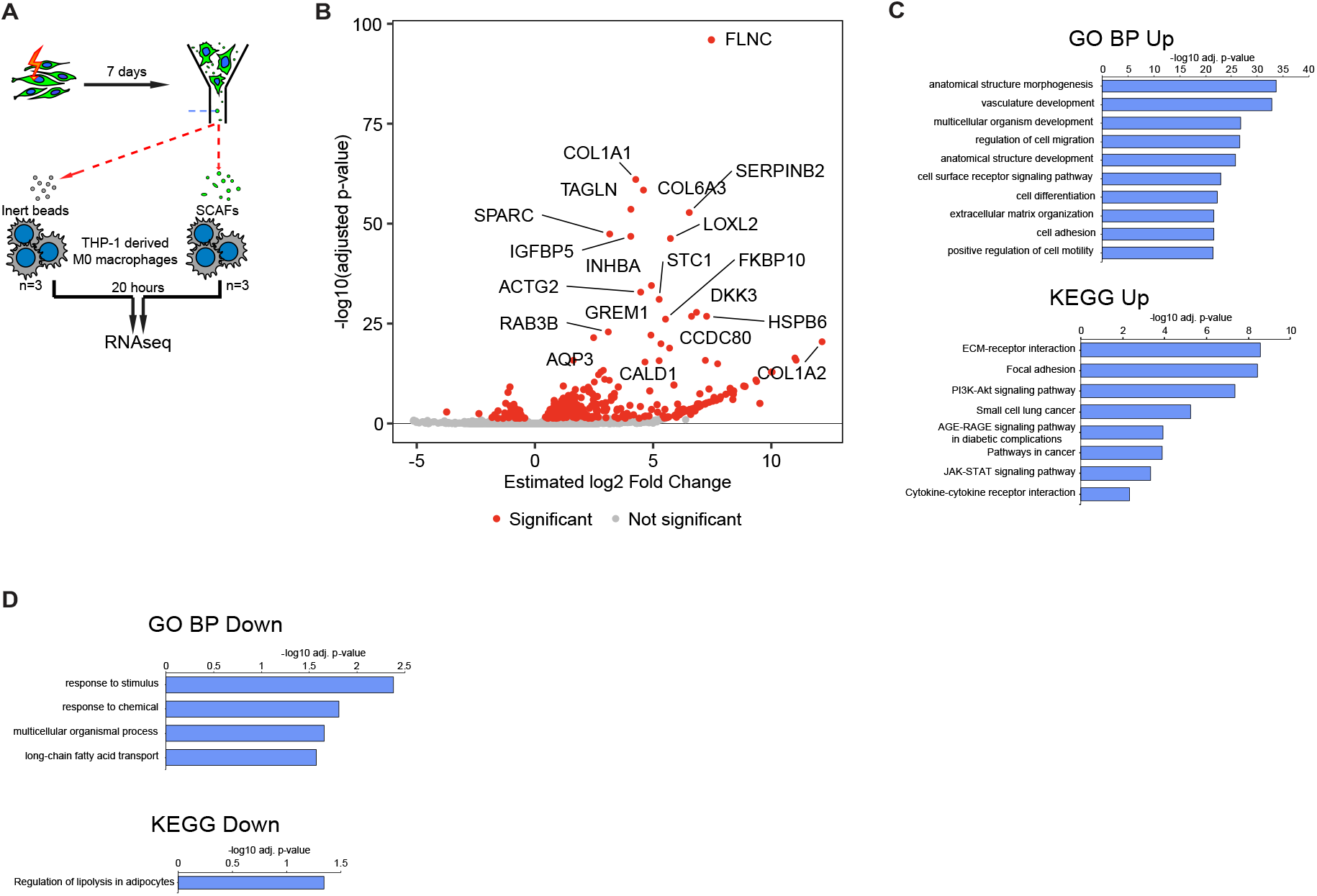
SCAF-treatment of THP-1 derived macrophages promotes cell migration and signaling pathways regulating macrophage activation. (A) Experimental design of RNA sequencing analysis of THP-1 derived macrophages after 20 hours of SCAF exposure. (B) Volcano plot of differentially expressed genes between macrophage cells treated with SCAFs compared to control cells treated with inert beads of similar size. (C) Bioinformatic analysis of significantly upregulated genes after SCAF-exposure using gProfiler. GO Biological process and KEGG pathway terms are shown. (D) Bioinformatic analysis of significantly downregulated genes after SCAF-exposure using gProfiler. GO Biological process and KEGG pathway terms are shown.

**Supplementary Table 1. SCAF Proteome**

List of the proteins that were identified in each of the replicate isolations of SCAFs that were analyzed by Mass Spectrometry.

